# Divergence of Grainy head affects chromatin accessibility, gene expression, and embryonic viability in *Drosophila melanogaster*

**DOI:** 10.1101/2024.04.07.588430

**Authors:** Henry A. Ertl, Erick X. Bayala, Mohammad A. Siddiq, Patricia J. Wittkopp

## Abstract

Pioneer factors are critical for gene regulation and development because they bind chromatin and make DNA more accessible for binding by other transcription factors. The pioneer factor Grainy head (Grh) is present across metazoans and has been shown to retain a role in epithelium development in fruit flies, nematodes, and mice despite extensive divergence in both amino acid sequence and length. Here, we investigate the evolution of Grh function by comparing the effects of the fly (*Drosophila melanogaster*) and worm (*Caenorhabditis elegans*) Grh orthologs on chromatin accessibility, gene expression, embryonic development, and viability in transgenic *D. melanogaster*. We found that the *Caenorhabditis elegans* ortholog rescued cuticle development but not full embryonic viability in *Drosophila melanogaster grh* null mutants. At the molecular level, the *C. elegans* ortholog only partially rescued chromatin accessibility and gene expression. Divergence in the disordered N-terminus of the Grh protein contributes to these differences in embryonic viability and molecular phenotypes. These data show how pioneer factors can diverge in sequence and function at the molecular level while retaining conserved developmental functions at the organismal level.

**SUMMARY STATEMENT:** Despite divergence in a disordered region that affects function at both molecular and organismal levels, the *Caenorhabditis elegans* Grainy head (Grh) protein rescued cuticle morphology in *D. melanogaster* embryos.

## INTRODUCTION

Metazoan development is driven by regulatory mechanisms that guide cellular growth and differentiation. Central to this mechanism are interactions between transcription factor proteins and *cis*-regulatory DNA that together control when, where, and how much a gene is expressed [1]. Some transcription factors, known as “master regulators”, play critical roles in major developmental transitions, such as driving pluripotency [2] and cell type differentiation [3]. Such master regulators are crucial for understanding the evolution of development because many of them are shared across metazoan life and have parallel developmental functions such as directing eye formation [4], appendage outgrowth [5], and epithelial cell differentiation [6,7]. However, despite this “deep homology” [8] – conservation in the underlying regulators of otherwise distinct morphologies (e.g., human and squid eye) – prior work has found that even deeply homologous master regulators have often diverged extensively in protein sequence and evolved distinct molecular functions [9–13].

To determine whether and how the molecular functions of shared master regulators have evolved, we must consider how these types of proteins act. As transcription factors, master regulators bind directly to *cis*-regulatory DNA and facilitate transcriptional activation and/or repression. Some master regulators — known as “pioneer factors” — have particularly widespread effects on gene regulation because of their unique ability to interact with nucleosome-bound DNA [7,14]. Before transcriptional activation can occur, such pioneer factors bind to nucleosome-bound DNA, evict nucleosomes, and allow for other regulatory factors to bind and modulate transcription [15,16]. Through this mechanism, pioneer factors can reshape the accessibility of the entire genome and drive cell type-specific regulatory programs [17,18]. Divergence in the sequence of a pioneer factor could therefore alter how and where it binds to DNA, how it affects chromatin structure, and ultimately how it impacts gene expression; changes in gene expression can then impact development.

Prior studies have found that the DNA binding specificity of transcription factors is generally well-conserved [19], which is consistent with greater levels of amino acid sequence conservation seen in the DNA binding domain relative to the rest of the protein [20]. Regions of transcription factors outside of the DNA binding domain are often unstructured and more rapidly evolving [20], but it has become increasingly appreciated that changes in these regions can also often affect how transcription factors affect regulation of downstream genes. For example, these regions can harbor important functions such as mediating protein-protein interactions [21], providing structural components needed to remodel chromatin [22], modifying DNA binding [23], and facilitating cooperative interactions with other transcriptional regulators [24,25]. For any particular transcription factor, the sequence-function relationship for regions outside of the DNA binding domain is often unclear, making it difficult to predict how evolutionary changes in this region will impact transcription factor function without empirical tests [26].

Here, we use such empirical tests to compare the functions of pioneer factor Grainy head (Grh) orthologs from *Drosophila melanogaster* and *Caenorhabditis elegans* during *D. melanogaster* embryogenesis. Grh is essential for epithelium development in flies, nematodes, and mice [6,27–29], and has been shown to act through pioneer factor mechanisms to “prime” *cis*-regulatory regions by making chromatin accessible at these regions [7,30]. Using transgenic fly strains expressing each of these orthologous proteins, we find functional conservation in the role of Grh in embryonic morphology but divergence in its functions affecting embryonic viability. We then characterize molecular phenotypes that capture multiple levels of the Grh regulatory mechanism by combining previously reported data on Grh binding with new data describing chromatin accessibility (ATAC-seq) and mRNA abundance (RNA-seq). Comparing these data between transgenic fly strains carrying the different Grh orthologs revealed differences in their molecular functions. Chimeras between the *D. melanogaster* and *C. elegans* Grh proteins as well as targeted mutations to the *D. melanogaster* allele identified specific regions in the N-terminal region of the protein potentially responsible for this functional divergence. These data underscore the importance of protein sequence changes outside of DNA binding domains and show that key developmental functions of pioneer factors can be maintained despite changes in protein sequence and molecular activities in distantly related organisms separated by 500 million years of evolution.

## RESULTS

### Evolution of Grh protein structure across metazoa

The *Grainy head* (*Grh*) gene was first identified and named in *Drosophila melanogaster* based on the large effects of loss-of-function mutations on the structure and melanization of larval mouth hooks that gave the appearance of a “grainy head”. Since then, *Grh* orthologs have been found in many metazoan species, where they have a conserved role in driving the differentiation of epithelial cells [28]. Canonical Grh isoforms expressed in the epidermis of *D. melanogaster* (fly-Grh) and *Caenorhabditis elegans* (worm-Grh) have relatively well-conserved DNA binding domains **(Figure 1A, B)**, consistent with prior work showing these orthologs bind to identical motifs [31]. However, the fly-Grh protein has a substantially longer disordered region than the worm-Grh protein on the N-terminus and contains an isoleucine-rich transactivation domain [32] **(Figure 1A)**. Comparison of Grh protein structure across metazoan orthologs **(Figure 1C)** suggests that the N-terminus expanded before the divergence of Ecdysozoa and Lophotrochozoa with a subsequent contraction along the Nematoda lineage. Within the expanded N-terminus, an isoleucine-rich transactivation domain (previously described by [32]) seems to have evolved along the Diptera lineage **(Figure 1C)**. The Grh protein, despite its well-conserved DNA binding domain, therefore seems to have evolved in other aspects of its structure. How these changes in the fly-Grh ortholog affect molecular and developmental processes remains unknown despite the highly conserved developmental role and DNA binding functionality of Grh proteins.

**Figure 1.**
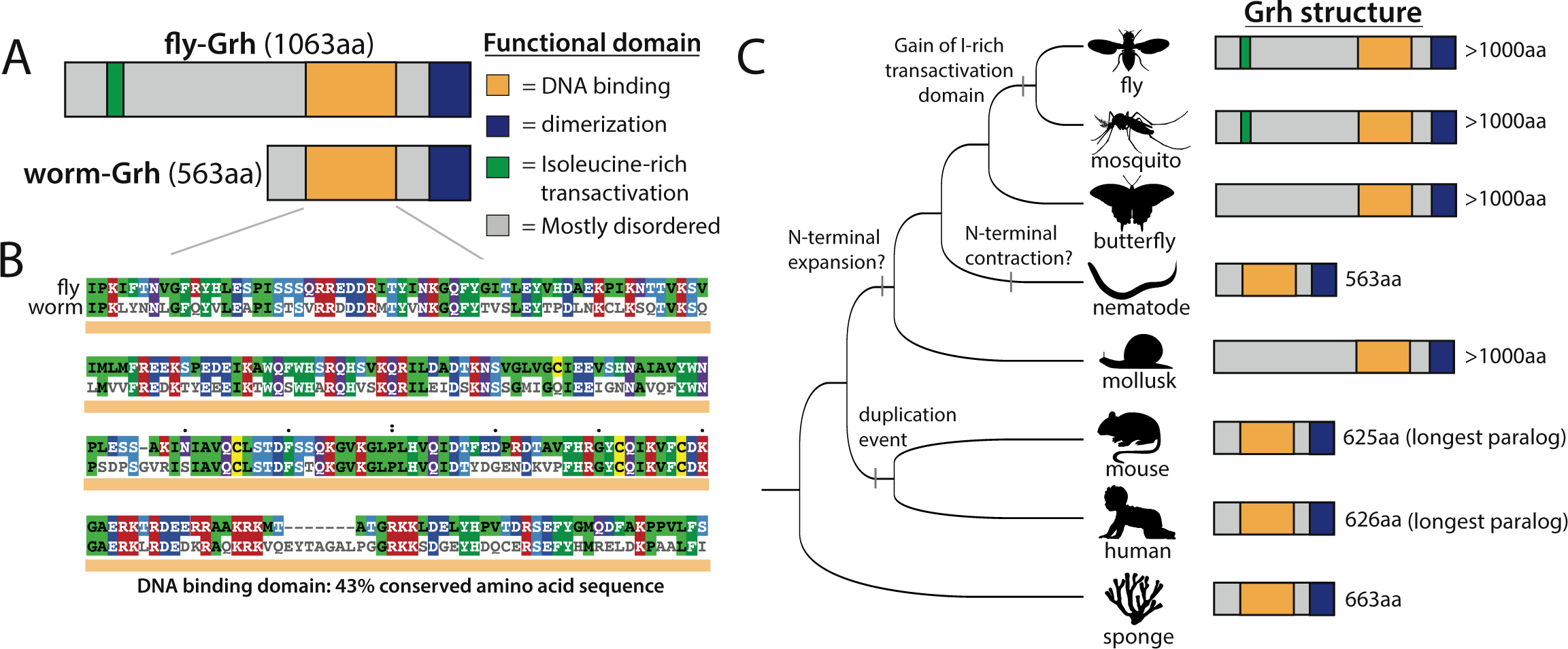
The structure of Grainy head across metazoa. **(A)** Schematic of *D. melanogaster* and *C. elegans* Grh (fly-Grh and worm-Grh, respectively) including color-coded functional domains. **(B)** Amino acid alignment of fly-Grh (top) and worm-Grh (bottom) annotated DNA binding domains. **(C)** Phylogeny of sampled metazoan species with schematics of Grh structure, similar to (A). Structure and lengths determined by longest isoform and paralog.

### *C. elegans* Grh rescues embryonic cuticle integrity in *D. melanogaster grh mutants*

To determine whether the molecular functions of Grh orthologs have diverged, we compared the functions of fly- and worm-Grh in the *D. melanogaster* embryo. More specifically, we constructed transgenic flies (*UAS-grh*) in a *grh* null mutant background that, when crossed with an activator line (*grh-Gal4*), express the *grh* transgenes of choice in a similar pattern to endogenous Grh during *D. melanogaster* embryogenesis. We created these flies by crossing each *UAS-grh* line, as well as a negative control “empty” UAS line, into a *grh* null mutant background (*grh^IM^*, [29]) and then crossing each of these lines to a *grh-Gal4* line that produces *gal4* transcripts in the *grh* expression pattern instead of functional *grh* transcripts [33] **(Figure S1A)**. Crossing the negative control UAS line in a *grh* null mutant background to this *grh-Gal4* line resulted in embryonic lethality and a near-complete loss of *grh* mRNA signal after the early stop codon contained in the *grh^IM^* allele (**Figure S1B, C**). This transgenic expression system allowed us to isolate and compare the effects of fly- and worm-Grh on *D. melanogaster* development.

*D. melanogaster grh* null embryos exhibit a “blimp” phenotype: compared to wildtype, they are wider, shorter, and often have torn cuticle [34]. To test for functional divergence between fly- and worm-Grh in cuticle formation, we analyzed the integrity of cuticle preparations in *grh* null mutant embryos expressing either the fly- or worm-Grh proteins. We additionally characterized embryos with the empty UAS construct (hereafter referred to as *grh* nulls) and embryos heterozygous for *grh*, carrying one null (*grh^IM^*) mutant allele one wild-type *D. melanogaster grh* allele as controls. For each genotype, we quantified the cuticle integrity by tracing the outline of five cuticle preparations and calculating the aspect ratio (width/height) **(Figure 2A)**, reasoning that this metric should capture the blimp phenotype of shorter and wider embryos.

**Figure 2.**
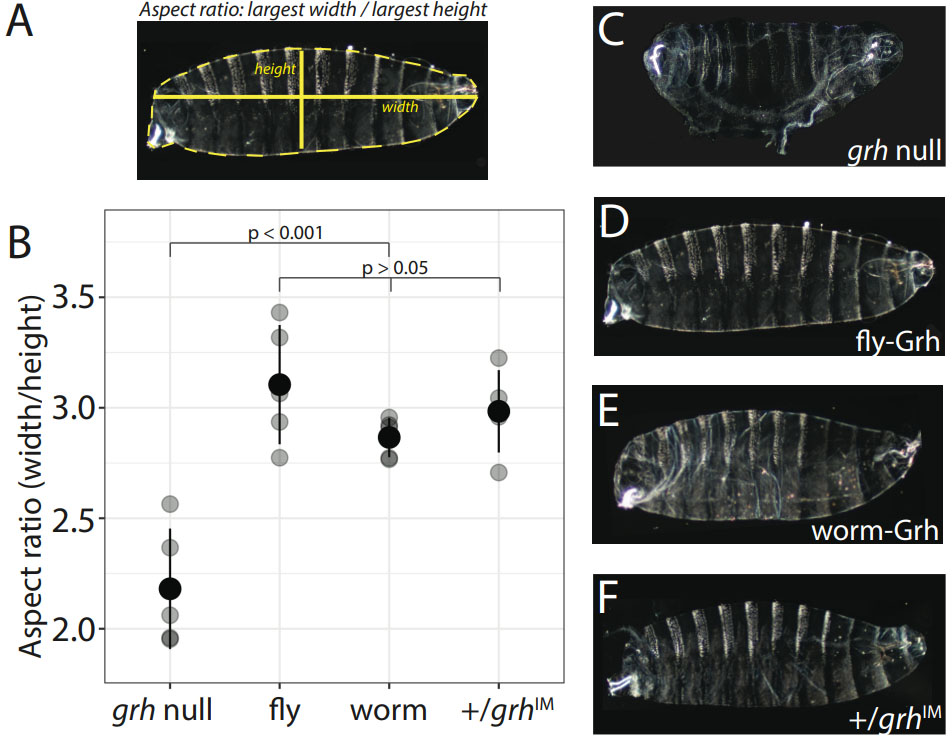
*C. elegans* Grainy head rescues embryonic cuticle integrity of *D. melanogaster Grh* mutants. **(A)** Example of how aspect ratio was measured by tracing the cuticle outline in ImageJ. **(B)** Aspect ratio for *grh* null embryos, embryos expressing fly-Grh or worm-Grh, and *grh* heterozygotes. Gray points show individual measurements, and black points and lines show the mean and standard deviation, respectively, of individual measurements for each genotype. See **Table S1** for all P values from Tukey post-hoc tests. **(C-F)** Cuticle preparations of unhatched first instar larvae expressing **(C)** no Grh transgene (i.e., *grh^IM^/grh^IM^ = “grh* null”), **(D)** fly-Grh, **(E)** worm-Grh, and **(F)** one wildtype *grh* allele (*grh*^+^/*grh^IM^ = “+/grh^IM^*”).

As expected, cuticle preparations of unhatched *grh* null embryos had consistent tearing in the cuticle and a significantly lower aspect ratio than embryos expressing the transgenic fly-Grh (p < 0.001, Anova with post-hoc Tukey, **Table S1**) (**Figure 2B, C and D, Figure S2**). Moreover, the aspect ratio of embryos expressing fly-Grh was not significantly different from that of embryos heterozygous for the recessive *grh* null allele (p = 0.97, Anova with post-hoc Tukey, **Table S1**) (**Figure 2B, D and F, Figure S2**), demonstrating that the *grh-Gal4* driver activated expression of the UAS transgenes in a manner sufficient to rescue endogenous Grh function in proper embryonic cuticle formation. Embryos expressing worm-Grh also had an aspect ratio that was not statistically significantly different from embryos expressing fly-Grh, suggesting that worm-Grh can fully rescue cuticle integrity (p = 0.56, Anova with post-hoc Tukey, **Table S1**) (**Figure 2B, D and E, Figure S2**). Taken together, these data suggest that substantial components of cuticle development function are conserved between the *D. melanogaster* and *C. elegans* Grh proteins.

### *C. elegans* Grh partially rescues embryonic viability of *D. melanogaster Grh* mutants

In addition to its role in cuticle formation, Grh also contributes to other developmental processes [35,36]. To compare the impacts of fly- and worm-Grh proteins on other embryonic development more generally, we quantified embryonic viability (i.e., hatching frequency) for *grh* null mutants expressing fly- or worm-Grh and compared these measures to the hatching frequency of sibling embryos that were either heterozygous or homozygous for the *grh^IM^* mutant allele. For each genotype, 6 replicates of 150-200 embryos each from the *UAS-grh* x *grh-Gal4* crosses were scored as hatched or unhatched at 30 hours after egg laying (AEL). None of the *grh* null embryos hatched, whereas their heterozygous siblings (which carried a wild-type *grh* allele) hatched ∼60% of the time (**Figure 3**). Embryos expressing fly-Grh hatched ∼39% of the time, indicating that our expression system restored a significant amount of Grh function during *D. melanogaster* embryogenesis (**Figure 3**, p < 0.001, Anova with post-hoc Tukey, **Table S2**). The difference in hatching frequency between embryos expressing fly-Grh controlled by the *grh-Gal4* driver and embryos heterozygous for a wildtype *grh* allele (39% vs 60%, respectively) (**Figure 3**, p < 0.001, Anova with post-hoc Tukey, **Table S2**) are presumably due to differences in expression level between the native *grh* gene and the *grh-Gal4* driver and/or the fact that the cDNA used in the UAS fly-Grh line produces only one Grh isoform. Embryos expressing worm-Grh in a *grh* null background hatched only ∼10% of the time, which was significantly lower than the ∼39% of embryos expressing the fly-Grh protein (**Figure 3**, p < 0.001, Anova with post-hoc Tukey, **Table S2**). Similar results were obtained using independent transgenic lines as well as transgenic lines in which a FLAG epitope (used for immunostaining to test for the proper nuclear subcellular localization of Grh) was fused to the C-terminal end of the Grh proteins (**Figure S3**). These data show that despite having similar effects on overall embryonic shape and general cuticle structure (**Figure 2**), the functions of the fly- and worm-Grh proteins have diverged in other ways that impact embryonic viability.

**Figure 3.**
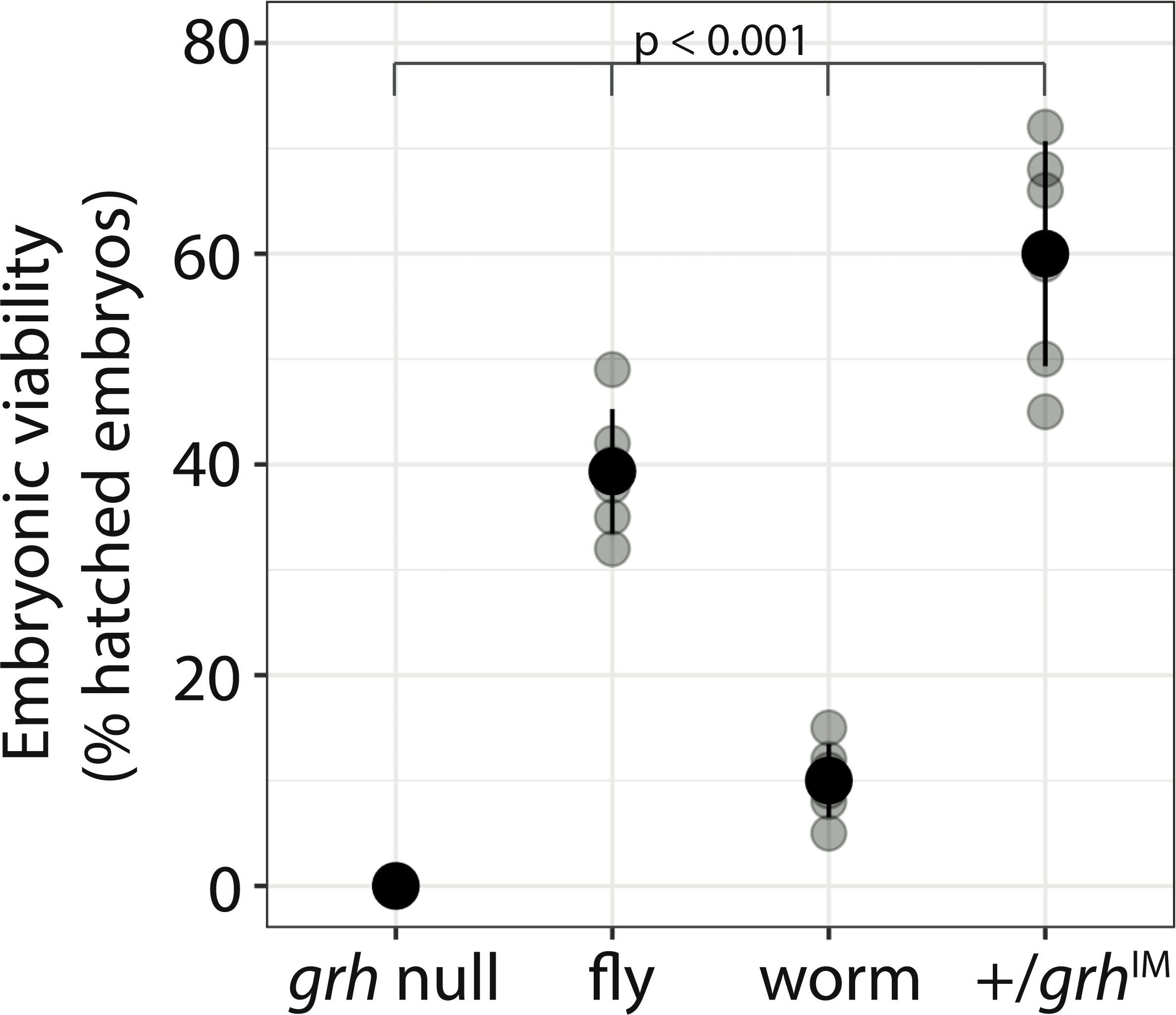
*C. elegans* Grh partially rescues embryonic viability of *D. melanogaster Grh* mutants. Embryonic viability, measured as percent hatched of 150-200 embryos, for *grh* null embryos, embryos expressing fly-Grh or worm-Grh, and *grh* heterozygotes (*grh^+^/grh^IM^*). Gray points show each measurement, and black points and lines show the mean and standard deviation, respectively, of the individual measurements. See **Table S2** for all P values from Tukey post-hoc tests.

### *C. elegans* Grh partially rescues chromatin accessibility at Grh-bound regions

Since Grh is a pioneer factor that, by definition, increases chromatin accessibility, we hypothesized that the failure of worm-Grh to fully rescue embryonic viability in *D. melanogaster* was likely due to divergence between the *D. melanogaster* and *C. elegans* Grh proteins that changed its impact on chromatin accessibility. To test this hypothesis, we collected ATAC-seq data from embryos 15-16 hours AEL, which is after the embryonic cuticle is formed and when Grh is regulating chromatin accessibility [36]. Because Grh is stably bound throughout embryogenesis [35], these data should also reflect differences in chromatin accessibility established by Grh earlier in development. We collected chromatin accessibility in embryos expressing the fly- or worm-Grh protein under the control of the *grh-Gal4* driver as well as from *grh* null embryos.

To first identify regions of chromatin with Grh-dependent accessibility in our transgenic system, we determined and compared accessible regions in *grh* null mutant embryos and embryos expressing fly-Grh. From this analysis (**Figure S4A**), we identified 31,446 regions with accessible chromatin in embryos expressing fly-Grh, but only 95 of these regions showed evidence of differential chromatin accessibility when compared with *grh* null embryos (**Figure 4A**). This low number of Grh-dependent accessible regions is consistent with a prior study that also found that Grh pioneer factor activity affected less than one hundred regions during late *Drosophila* embryogenesis [36]. These 95 regions were enriched for canonical Grh binding motifs as well as for binding sites for Blimp-1, a transcription factor highly expressed in the epithelial cells that is also required for cuticle formation **(Figure 4B)**.

**Figure 4.**
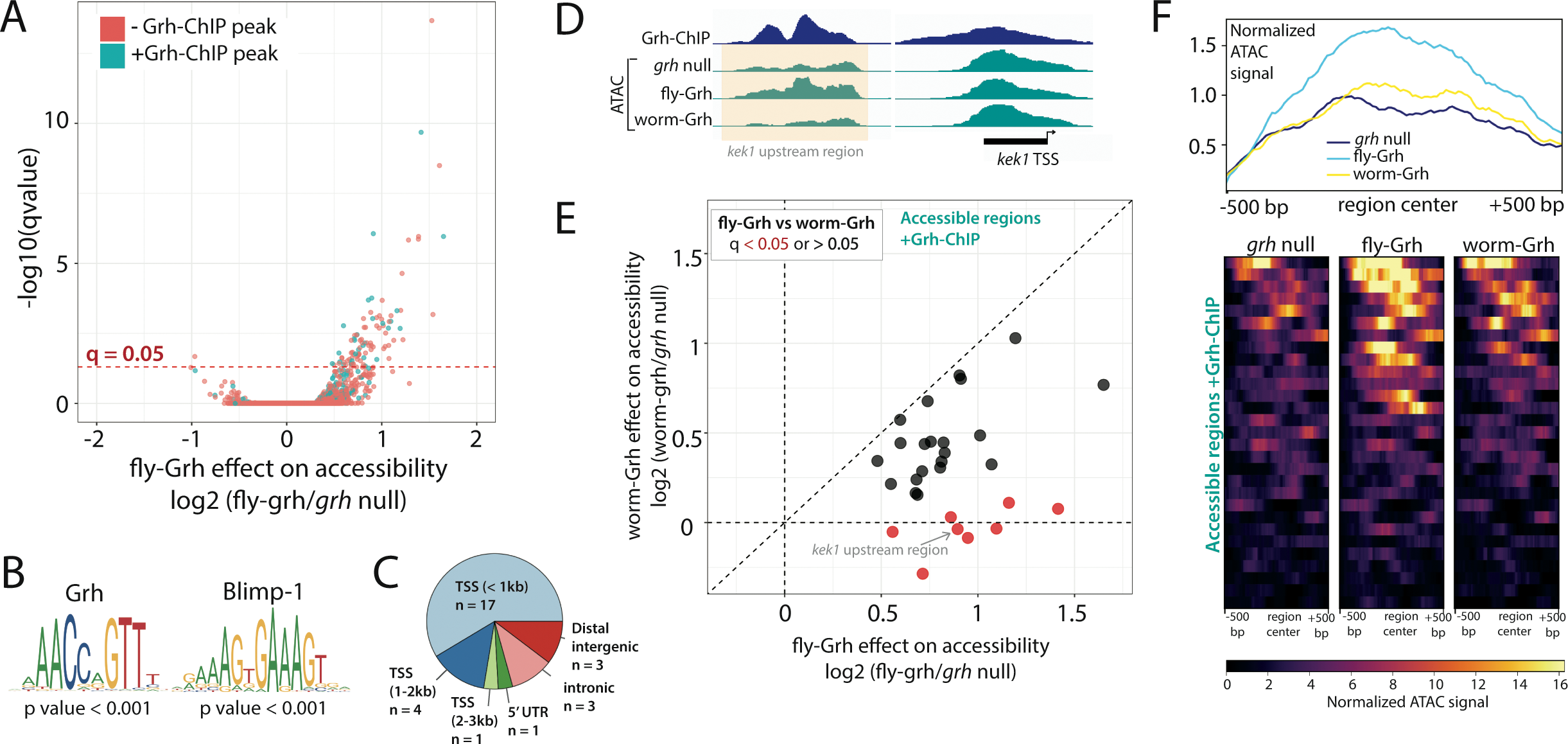
*C. elegans* Grh partially rescues chromatin accessibility of Grh-bound regions in *D. melanogaster.* **(A)** Volcano plot showing the relationship between the log2 fold change of chromatin accessibility between embryos expressing fly-Grh and *grh* null embryos versus -log10(q-value obtained from DEseq2). Regions are colored teal if they overlap with Grh ChIP-seq peaks from [35] and salmon if they do not overlap. **(B)** Position weight matrices that were significantly overrepresented (determined by *icistarget*) among all regions with q < 0.05 from (A). **(C)** Pie chart showing the numbers of regions falling into each genomic category. **(D)** Screenshot of genome browser showing the normalized ATAC signal for the region upstream of *kek1* (highlighted). Input-normalized ChIP-seq data from 15-16 hour AEL embryos collected by [35] is also shown. **(E)** Scatterplot contrasting the log2 fold change of chromatin accessibility between (x-axis) embryos expressing fly-Grh and *grh* null embryos and (y-axis) embryos expressing worm-Grh and *grh* null embryos. Points are color-coded to indicate whether regions have significantly different accessibility between embryos expressing fly- and worm-Grh (q < 0.05: red; q > 0.05: black). “Normalizing” by the *grh* null embryos distinguishes between Grh-dependent increases and decreases in accessibility. The *kek-1* upstream region highlighted in (D) is indicated with an arrow. **(F)** A metaplot is shown at the top summarizing the mean values from the heatmaps underneath displaying normalized ATAC signals of 1kb around the center of the 29 accessible regions for grh null embryos (black) and embryos carrying the fly-Grh (blue) or worm-Grh (yellow) transgenes.

To identify regions most likely to be directly affected by Grh, we compared these data to previously collected chromatin immunoprecipitation with sequencing (ChIP-seq) data for Grh in *D. melanogaster* embryos from the same 15-16 hours AEL timepoint [35]. We found that 29 of the 95 regions we identified as having Grh-dependent accessible chromatin were also reported to show evidence of Grh binding (**Figure 4A**, “+Grh-ChIP peak”, teal-colored points); other Grh-bound regions also showed differences in chromatin accessibility in our study but were not called statistically significant (**Figure 4A**). More than half of these 29 regions (17/29 = 59%) are located less than 1 kb from a transcription start site, with 22 of the 29 (76%) located within 3 kb of a transcription start site **(Figure 4C**). Among the rest, 3 are located in intronic sequences and 3 are located in intergenic regions further than 3 kb from a transcription start site **(Figure 4C**). An example of these data is shown for the *kek-1* gene, which encodes a transmembrane protein that is highly expressed in epithelial cells [37,38] (**Figure 4D**); note that the region ∼1 kb upstream of the *kek-1* transcription start site shows a reduction in accessibility in both *grh* null mutant embryos and embryos expressing worm-Grh when compared to embryos expressing fly-Grh.

All 29 of the Grh-dependent accessible regions with evidence of Grh binding in *D. melanogaster* showed a similar pattern to *kek-1*, with chromatin less accessible in embryos expressing worm-Grh than in embryos expressing fly-Grh (**Figure 4E**). Although only 8 of the 29 individual differences were statistically significant at the genomic level (FDR < 0.05, Benjamini Hochberg correction, **Figure 4E**), this pattern is unlikely to arise by chance (Wilcoxon sign test, p < 0.01). These differences in accessibility are not explained by global differences in chromatin accessibility in embryos expressing worm-Grh and fly-Grh: a strong correlation (r = 0.99) in chromatin accessibility was observed for all 31,446 accessible regions (**Figure S4**). The differences observed between embryos expressing fly- and worm-Grh are also unlikely to be explained by a complete lack of chromatin remodeling function for the worm-Grh protein because chromatin accessibility in embryos expressing worm-Grh was generally greater in these 29 regions than chromatin accessibility in *grh* null mutants (**Figure 4F**). We found only two regions of the genome with significant differences in chromatin accessibility between *grh* null embryos and embryos expressing worm-Grh but not fly-Grh. This result suggests that the worm-Grh protein can, but does not often, regulate chromatin accessibility at places in the genome that fly-Grh does not (**Figure S5A**). Taken together, these data suggest that divergence between the *C. elegans* and *D. melanogaster* Grh proteins affects the extent of chromatin accessibility in *D. melanogaster* embryos more than it affects where it binds, consistent with prior work suggesting these two proteins bind to the same sites [31].

### *C. elegans* Grh under-activates Grh direct targets

Given that a primary function of pioneer factors is to make chromatin accessible for other transcription factors, we hypothesized that the reduced chromatin accessibility seen in embryos expressing worm-Grh rather than fly-Grh could have downstream consequences on gene expression. Furthermore, and perhaps independent of the effects of protein sequence divergence on chromatin accessibility, we also reasoned that the presence of an isoleucine-rich domain with transactivation abilities in fly-Grh that is absent in worm-Grh might also cause these orthologous proteins to have different effects on gene expression. To test these possibilities, we used RNA-seq to measure mRNA abundance in embryos expressing fly-Grh, embryos expressing worm-Grh, and *grh* null mutant embryos, and then we compared the expression levels of putative Grh direct targets identified in a previous study [35].

First, we sought to confirm that the previously identified direct targets of *D. melanogaster* Grh showed evidence of Grh regulation in our UAS/Gal4 system by comparing RNA-seq data from embryos expressing fly-Grh to RNA-seq data from *grh* null mutant embryos. We found that 415 of the 1614 previously identified Grh direct targets were significantly differentially expressed (FDR < 0.05, Benjamini Hochberg correction), with 276 genes being up-regulated by fly-Grh and 139 genes being down-regulated (**Figure 5A, Figure S6A**), consistent with prior data showing that Grh can have positive or negative effects on transcript levels [6,35]. Of these 415 genes, a select GO enrichment analysis [39] showed (**Figure 5B**) that (1) proteins with DNA-binding activity are underrepresented, (2) proteins with structural cuticle activity are enriched, and (3) proteins with oxidoreductase activity are also enriched. These results suggest that Grh is not upstream of many regulatory transcription factors and are consistent with the well-documented role of Grh in cuticle formation [34]The relationship between Grh and regulation of oxidoreductase enzymes remains unknown.

**Figure 5.**
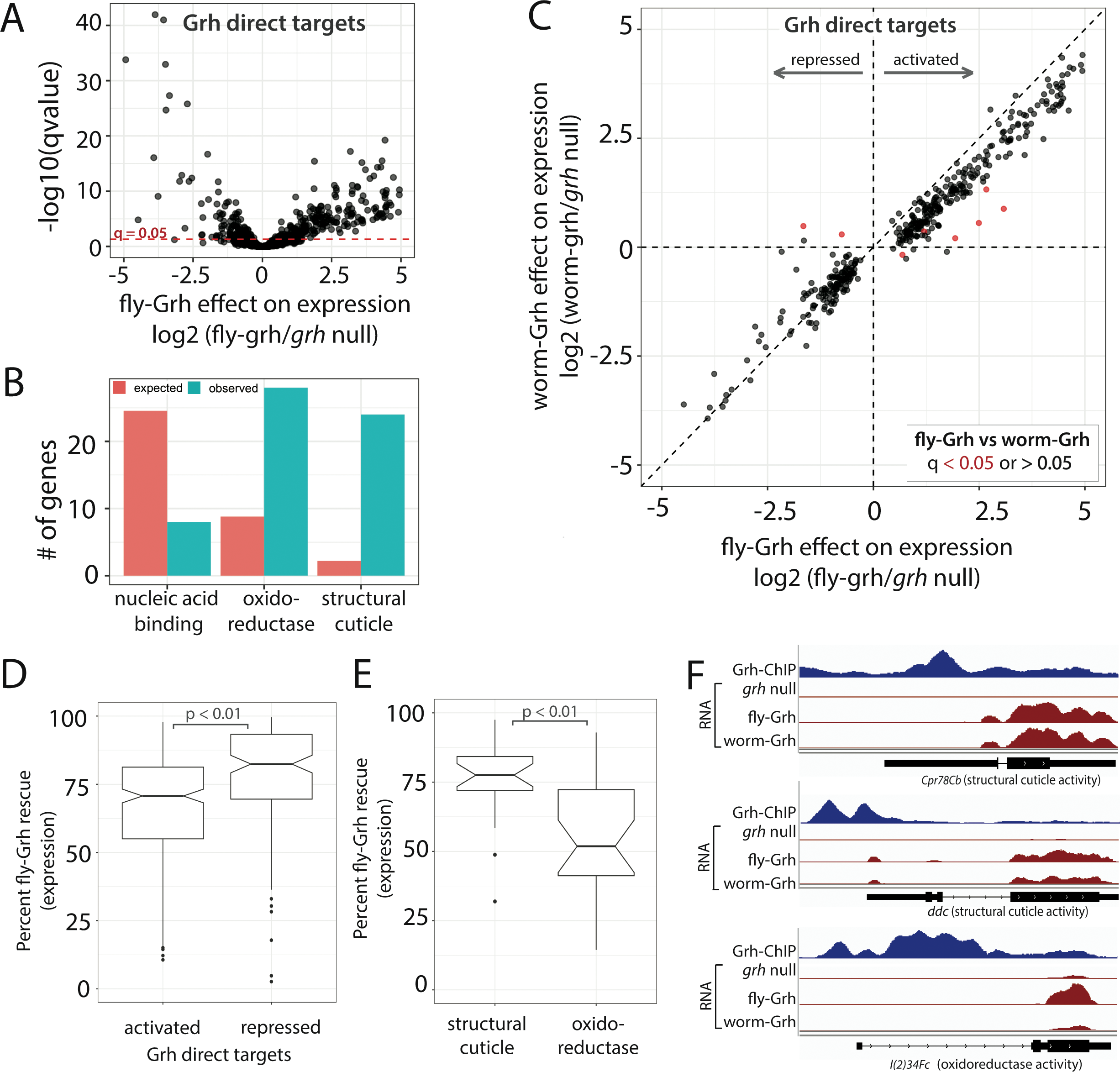
*C. elegans* Grh under-activates *D. melanogaster* Grh direct targets. **(A)** Volcano plot showing the relationship between the log2 fold change of gene expression between embryos expressing fly-Grh and *grh* null embryos versus -log10(q-value obtained from DEseq2). Genes with q-values < 0.05 are classified as fly-Grh direct targets and used for subsequent analyses. **(B)** Expected (salmon) and observed (teal) numbers of genes for significantly enriched or depleted GO slim molecular categories from the list of fly-Grh direct targets. **(C)** Scatterplot contrasting the log2 fold change of gene expression between (x-axis) embryos expressing fly-Grh and *grh* null embryos and (y-axis) embryos expressing worm-Grh and *grh* null embryos. “Normalizing” by the *grh* null embryos distinguishes between Grh-dependent activation and repression. Points are color-coded to indicate whether regions have significantly different expression levels between embryos expressing fly- and worm-Grh (q < 0.05: red; q > 0.05: black). **(D)** Box plots contrasting the percent fly-Grh rescue in embryos expressing worm-Grh for Grh-activated (log2(fly-Grh/*grh* null) > 0) and Grh-repressed (log2(fly-Grh/*grh* null) < 0) genes. Notches represent 95% confidence intervals around the median. P-value from Wilcoxon paired rank sum test. **(E)** Box plots contrasting the percent fly-Grh rescue in embryos expressing worm-Grh for fly-Grh direct target genes with GO slim molecular categories “structural cuticle” and “oxidoreductase”. Notches represent 95% confidence intervals around the median. P-value from Wilcoxon paired rank sum test. **(F)** Screenshots from genome browser show RNA-seq read counts normalized as counts per million (CPM) from *grh* null embryos and embryos expressing either fly-Grh or worm-Grh for the *Cpr78Cb*, *ddc*, and *l(2)34Fc* genes. Input-normalized ChIP-seq data from 15-16 hour AEL embryos collected by [35] is also shown for each gene.

We next tested the extent to which the worm-Grh protein rescues expression of these 415 genes by comparing gene expression in embryos expressing the worm- or fly-Grh proteins to gene expression in *grh* null embryos **(Figure 5C).** We found that up-regulated genes are consistently under-activated in embryos expressing worm-Grh; by contrast, the difference between worm-Grh and fly-Grh is less pronounced for down-regulated genes (p < 0.01, Wilcoxon rank sum test) **(Figure 5D)**. This significant pattern was observed even though only 19 genes showed a statistically significant difference at a genomic level between embryos expressing fly-Grh versus worm-Grh (FDR < 0.05, Benjamini Hochberg correction, **Figure 5C**). We note that despite the skewed expression of these Grh target genes, the overall pattern of gene expression was similar between embryos expressing the fly- and worm-Grh proteins (r = 0.98, **Figure S6B**). We also note that 34 genes showed evidence of differential expression between *grh* null embryos and embryos expressing worm-Grh but not fly-Grh, suggesting that the *C. elegans* Grh protein might regulate a few different genes in *D. melanogaster* embryos than the *D. melanogaster* Grh protein (**Figure S5B**).

Finally, we looked at the impacts of worm-Grh on expression of the sets of genes annotated as structure cuticle or oxidoreductase activity, which were enriched for expression differences between embryos expressing fly-Grh and *grh* null embryos. For each of these genes, we calculated the “percent fly-Grh rescue” (see Methods) using the RNA-seq data from embryos expressing worm-Grh. We found that the 24 structural cuticle genes were expressed at an average of 77% of the levels seen in embryos expressing fly-Grh (**Figure 5E**). By contrast, we found that the 28 oxidoreductase genes were expressed at an average of only 51% of the levels seen in embryos expressing fly-Grh **(Figure 5E**), which was a significantly greater reduction in expression than seen for the structural cuticle genes (p < 0.01, Wilcoxon rank sum test). Three specific examples of genes from these functional groups are shown in **Figure 5F**: the worm-Grh protein caused *Cpr78Cb* and *ddc*, both of which are involved in cuticle development, to be expressed at 95% and 87%, respectively, of the levels activated by fly-Grh, whereas it caused the oxidoreductase gene highly expressed in epithelial cells, *l(2)34Fc*, to be expressed at only 29% of the levels activated by fly-Grh. These observations are consistent with – and potentially provide an explanation for – our finding that worm-Grh fully rescues cuticular morphology but not embryonic viability.

### Differences in the N-terminus of Grh contribute to functional divergence between the *C. elegans* and *D. melanogaster* orthologs

To determine which part(s) of the divergent worm- and fly-Grh proteins were responsible for their differences in ability to rescue embryonic viability of *grh* null mutants as well as their effects on chromatin accessibility and gene expression, we constructed and analyzed chimeric Grh proteins expressed using the same UAS/Gal4 system used to analyze the species-specific Grh alleles. Because our data as well as prior work [31] indicate that worm- and fly-Grh bind to identical sequences, and because there is substantial sequence and length variation outside of the DNA binding domain, we reasoned that at least some of the functional differences we observe between worm- and fly-Grh proteins might be due to changes in the largely disordered N-terminal region. To test this hypothesis, we examined the functional effects of chimeric proteins in which the N-terminus (defined here as ending 10 amino acids upstream of the DNA binding domain) was reciprocally swapped between fly- and worm-Grh (hereafter referred to as worm/fly-Grh and fly/worm-Grh) (**Figure 6A**). In addition, to specifically determine the effects of the isoleucine-rich transactivation domain present in fly-Grh but absent in worm-Grh, we created a fly-Grh transgene in which each isoleucine in this domain was changed to an alanine (hereafter referred to as fly-ItoA-Grh) [32] (**Figure 6A**). For all three of these modified proteins, two independent transgenic UAS lines were created and analyzed, showing similar results when activated by the *grh-Gal4* driver (**Figure S3A**).

**Figure 6.**
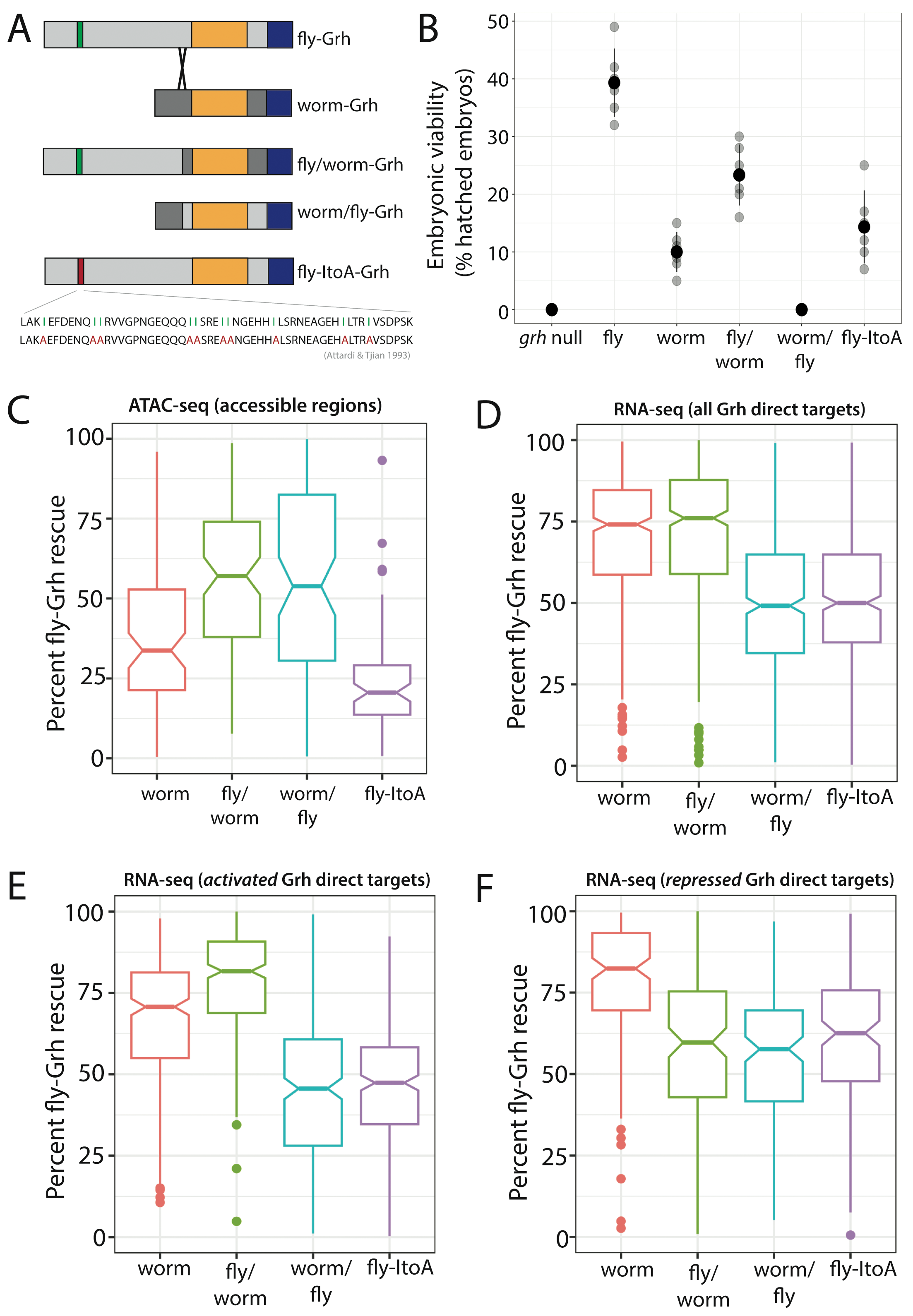
Functional divergence between Grh orthologs can be partially explained by differences in the N-terminus. **(A)** Schematic of all chimeric and mutant Grh transgenes. Functional domains are color-coded as in Figure 1A. The sequence changes in the isoleucine-rich transactivation domain from fly-Grh to fly-ItoA-Grh are shown with amino acid differences highlighted in green and red, respectively. **(B)** Embryonic viability, measured as percent hatched of 150-200 embryos, for *grh* null embryos and embryos expressing fly-Grh, worm-Grh, fly/worm-Grh, worm/fly-Grh and fly-ItoA-Grh. Gray points = individual measurements, and black points and lines show the mean and standard deviation, respectively, of the individual measurements for that genotype. **(C)** Percent fly-Grh rescue of chromatin accessibility for 29 regions in all transgenic strains. See Table S3A for all P values from Tukey post-hoc tests. **(D-F)** Same as C but for expression of all 415 Grh-regulated genes **(D)**, as well as the subsets of 276 activated **(E)** and **(F)** 139 repressed Grh direct target genes. Genes are defined as activated or repressed if their expression = log2(fly-Grh/*grh* null) > OR < 0). See Table S3B for all P values from Tukey post-hoc tests.

For each of these modified Grh proteins, we measured their effects on embryonic viability using the same UAS/Gal4 expression system described above and compared their effects to those of the intact fly- and worm-Grh proteins. We found that embryos expressing fly/worm-Grh (i.e. Grh with the N-terminus from fly-Grh and the C-terminus from worm-Grh) have significantly higher embryonic viability than embryos expressing worm-Grh (23% and 10%, respectively) (p < 0.001, Anova with post-hoc Tukey) **(Figure 6B**), consistent with the hypothesis that sequence divergence in the N-terminal region contributes to functional divergence between worm- and fly-Grh proteins. Interestingly, embryos expressing worm/fly-Grh (i.e. Grh with the N-terminus from worm-Grh and the C-terminus from fly-Grh) did not hatch at all (**Figure 6B**), showing significantly lower embryonic viability than embryos expressing the intact worm-Grh protein (0% versus 10% embryonic viability, respectively) (p < 0.001, Anova with post-hoc Tukey). This observation suggests that there is an epistatic interaction between divergent sites in the N-terminal and C-terminal regions of Grh. Embryos expressing fly-ItoA-Grh (i.e. fly-Grh with isoleucines changed to alanines in the transactivation domain) hatched 14% of the time on average (**Figure 6B**), which was significantly lower than embryos expressing fly-Grh (p < 0.001, Anova with post-hoc Tukey), confirming a functional role for the isoleucine-rich domain in fly-Grh that is absent in worm-Grh.

Next, we measured the effects of these chimeric fly/worm-Grh and worm/fly-Grh proteins as well as the fly-ItoA-Grh protein on chromatin accessibility using ATAC-seq, as described above for embryos expressing the fly-Grh and worm-Grh proteins and for *grh* null mutant embryos. For each of the modified Grh proteins, we measured their effects relative to embryos expressing fly-Grh by calculating the “percent fly-Grh rescue”. In the 29 regions described above as showing Grh-dependent chromatin accessibility in *D. melanogaster*, we found that embryos with expression of the fly/worm-Grh or worm/fly-Grh protein showed increased chromatin accessibility relative to that seen in embryos expressing worm-Grh (**Figure 6C**, **Figure S7A-C**, **Table S3A**). This observation indicates that divergent sequences in both the N-terminal and C-terminal portions of the *D. melanogaster* Grh protein increase its ability to make chromatin more accessible, however the isoleucine-rich region in the N-terminus seems to be particularly important because embryos expressing the fly-ItoA-Grh protein showed less chromatin accessibility in these regions than embryos expressing worm-Grh (**Figure 6C**, **Figure S7A and D**, **Table S3A**).

For gene expression, we measured the “percent fly-Grh rescue” seen in RNA-seq data collected from embryos expressing these modified Grh proteins. When considering all 415 Grh-regulated genes, we found that the median expression level restored by the worm-Grh and fly/worm-Grh proteins was about 75% whereas the median expression level restored by the worm/fly-Grh and fly-ItoA proteins was closer to 50% **(Figure 6D, S7E-H)**. When we analyzed the 276 Grh-activated and 139 Grh-repressed genes separately, however, we observed that the fly/worm-Grh protein caused expression of Grh-activated genes to be most similar to embryos expressing fly-Grh whereas the worm-Grh protein had this effect for Grh-repressed genes (**Figure 6E and F, Table S3**). Embryos expressing the worm/fly-Grh protein restored even less expression of Grh-activated genes in *grh* null embryos than the worm-Grh protein (**Figure 6E**, **Table S3**). Finally, embryos expressing the mutant fly-ItoA-Grh protein showed expression most similar to embryos expressing the worm/fly-Grh protein for both Grh-activated and Grh-repressed genes (**Figure 6E and F, Table S3**). Taken together, these observations suggest that the isoleucine-rich domain in the N-terminus that was gained in the lineage leading to *D. melanogaster* (**Figure 1C**) contributes more to Grh-dependent activation than repression of gene expression in *D. melanogaster* embryos.

Considering the effects of the chimeric and mutant Grh proteins on chromatin accessibility, activation of gene expression, and embryonic viability together suggests that their effects are more complex than a simple model in which Grh binds to chromatin, makes the DNA more accessible, and activates gene expression, resulting in proper development that produces viable embryos. For example, we found that combining the N-terminal (transactivation) region of fly-Grh with the C-terminal (DNA-binding) region of worm-Grh increased chromatin accessibility, expression activation, and embryonic viability (**Figure 6B, D, and E**), but even though similar chromatin accessibility was seen when combining the N-terminal region of worm-Grh with the C-terminal region of fly-Grh (**Figure 6C**), we observed reduced activation of gene expression (**Figure 6E**) and the complete loss of embryonic viability also seen in *grh* null mutants (**Figure 6B**). Furthermore, the fly-ItoA-Grh protein, in which isoleucines in the N-terminal region of the *D. melanogaster* protein were changed to alanines (**Figure 6A**), resulted in less chromatin accessibility than the much more divergent *C. elegans* Grh protein (**Figure 6C**) and reduced expression activation similar to the worm/fy-Grh chimeric protein (**Figure 6E**), yet resulted in greater embryonic viability than the worm-Grh protein (**Figure 6B**). These observations underscore the need for further work elucidating the functional connections between chromatin structure, gene expression, and development.

## DISCUSSION

In this study, we compared the molecular and developmental functions of orthologous Grh proteins from *D. melanogaster* and *C. elegans* in *D. melanogaster* embryos. We found that these two highly divergent proteins appear to still bind to similar sequences in the genome (consistent with prior work, e.g., [19], but they increase chromatin accessibility and gene expression to different extents. This difference in molecular impacts of the fly- and worm-Grh proteins seems primarily attributable to differences in the N-terminal region, particularly in the derived isoleucine-rich transactivation domain that arose in the Diptera lineage. Despite these differences in molecular functions, the *D. melanogaster* and *C. elegans* Grh proteins retain similar effects on the development of embryonic cuticle in *D. melanogaster* but result in different degrees of embryonic viability, which might be due to the impact of Grh on expression of genes with oxidoreductase activity. Taken together, these data show that the binding of Grh is distinct from its ability to increase chromatin accessibility, which is at least somewhat distinct from its ability to activate gene expression 一 and its effects on the regulation of gene expression do not always correlate with its impacts on organismal traits like embryonic cuticle structure and embryonic viability.

Despite the common narrative that transcription factor functions are largely conserved over long evolutionary timescales, which is supported by deeply conserved amino acid sequences of DNA binding domains [19,20], our work comparing the effects of fly-Grh and worm-Grh in *D. melanogaster* embryos, as well as other experimental tests comparing activities of divergent transcription factors in transgenic animals (**Table S4**), paints a different picture. Even early empirical case studies that emphasized a surprising conservation of function, such as the mouse eyeless/Pax6 protein driving development of ectopic eye tissue in *D. melanogaster* [4], were later suggested to have functionally diverged with the *D. melanogaster* Eyeless protein inferred to possibly have evolved a novel repressor domain not found in the mouse ortholog [40]. Similarly, when the fly ortholog of Engrailed was expressed in the mouse, it rescued loss-of-function phenotypes in the mid-hindbrain but did not rescue limb development defects [9]. These types of observations have led some to argue that the role of variation in transcription factor function in gene expression evolution might be under-appreciated [20,41,42]. Detailed studies of such divergent functions have been sparse, however, because most of these functional comparisons between divergent transcription factor orthologs were done between 1996 and 2002 (**Table S4**) when tools for studying the molecular functions of a transcription factor on a genomic scale were much more limited.

With strong conservation typically seen for DNA binding domains, much of the functional divergence of transcription factors might be due to changes in other regions of the protein. Indeed, these regions, which can contain functional domains responsible for dimerization, transactivation, and post-translational modification, for example, are often much less conserved than DNA binding domains and have been shown to contribute to major evolutionary innovations [43]. Many transcription factors have disordered regions outside of their DNA binding domains that might have particularly few sequence constraints, enabling their evolutionary divergence. While it is possible – if not likely – that a significant amount of amino acid sequence variation in such regions is non-functional, recent work has shown that these disordered regions can influence DNA binding, mediate protein-protein interactions, and affect phase separation [23,44]. Our work on Grainy head supports these ideas in that we found sequence divergence between flies (*D. melanogaster*) and worms (*C. elegans*) in the N-terminal, largely disordered region of the protein, that affects its ability to make chromatin accessible, activate gene expression, and maintain viability of *D. melanogaster* embryos. The evolution of one particular portion of the *D. melanogaster* N-terminal region (i.e., the isoleucine-rich domain) was found to be primarily responsible for these divergent functions.

Most transcription factors bind to accessible *cis*-regulatory DNA and affect transcription, but some transcription factors (including Grh) act as pioneer factors that also influence gene expression by altering chromatin structure. Since chromatin accessibility is both an effect and effector of transcription factor binding [45], it is often assumed that variation in chromatin accessibility alters gene expression; however, work in a variety of systems suggests that this relationship is not so straightforward. For example, in *Saccharomyces species*, variation in chromatin accessibility does not correlate with that of gene expression [46], yet population variation in chromatin accessibility collected from *Drosophila* embryos correlates more strongly with variation in gene expression, with the strongest correlation seen for regions close to a transcription start site [47]. Important questions remain about the relationship between chromatin accessibility, transcription factor binding, and gene expression [48–51], and the work described here comparing the molecular effects of divergent fly-Grh and worm-Grh in *D. melanogaster* embryos shows how protein divergence can have different effects on each of these levels.

In summary, both prior work and the work presented here show that transcription factor evolution involves both deep functional conservation *and* divergence. Some functions of transcription factors – most often DNA binding – are conserved over hundreds of millions of years, while others, including impacts on chromatin accessibility and gene expression, evolve more rapidly and can contribute to the evolution of gene expression and development. Some developmental functions (e.g., embryonic cuticle formation regulated by Grh) appear to be less sensitive to some divergence in molecular function than others (e.g., Grh’s effects on embryonic viability). More functional work in transgenic animals is needed to gain a more comprehensive understanding of transcription factors and their effects on gene regulatory and developmental evolution.

## MATERIALS & METHODS

### Fly stocks

The following used lines were from the Bloomington Drosophila Stock Center: y[1], w[1118]; PBac{y[+]-attP=3B}VK00037 (#9752), w[*]; sna[Sco]/CyO, P{Dfd-GMR-nvYFP}2 (#23230), and y[1], w[*]; Mi{Trojan-GAL4.0}grh[MI05134-TG4.0]/SM6a (#76172). The *grh^IM^* line was a gift from Bill McGinnis.

### Molecular cloning

pUAST-attB and pUAST-attB-3xFLAG vectors were PCR linearized (**Table S5**) and purified using NEB PCR and DNA cleanup kit. The coding sequences of the canonical isoforms of *grh* expressed in the epidermis of *D. melanogaster* and *C. elegans* (for fly- and worm-Grh, respectively) were generated by *de novo* synthesis (IDT), and then amplified with overhangs corresponding to the linearized regions pUAST-attB and pUAST-attB-3xFLAG vectors and purified. fly- and worm-Grh fragments were each cloned into linearized pUAST-attB and pUAST-attB-3xFLAG vectors using Gibson Assembly [52]. For the chimeric transgenes (fly/worm-Grh and worm/fly-Grh), the chimeric fragments were amplified off of already-constructed fly- and worm-Grh plasmids and cloned together with Gibson Assembly. For the mutated fly-Grh transgene (fly-ItoA-Grh), the seven isoleucines were changed to alanines using site-directed mutagenesis. For all PCR primers used in molecular cloning, see **Table S5**. All final plasmids were sequenced using long-read sequencing (Plasmidsaurus) and the resultant plasmid maps are provided in the supplementary information.

### Drosophila transgenesis

The VK37 line was selected for use with the ϕC31-attP transgenesis system because prior work found that Gal4 drove a UAS-Grh transgene similarly to endogenous Grh expression [29]. The VK37 landing site was recombined onto a chromosome with the in *grh*^IM^ allele, using the *yellow^+^* (*y^+^*) marker to select for the landing site and Sanger sequencing to verify the *grh^IM^* allele. This line containing the VK37 landing site and the *grh^IM^* allele was injected by Genetivision with each of the UAS plasmids described above. F1 transformants that carried the *white*^+^ (*w^+^*) allele, indicating successful transformation of the UAS construct, were then balanced with a CyO chromosome that expresses green fluorescent protein in embryos (#23230) in a *yw* background. Transgenic lines were tested for correct Grh transformation with PCR. Two independent transformation strains were maintained for each Grh transgene.

### UAS-Grh x Grh-Gal4 cross and embryo collections

To generate the *Grh-Gal4* line for the main experimental cross, the *Grh-Gal4* allele (#76172) line was similarly crossed into a *yw* background and balanced with the same fluorescent CyO chromosome (#23230) as the transgenic lines described above. For one cross, ∼500 virgin females of this *Grh-Gal4* line were collected over five days and then combined with ∼30 young males of a UAS-Grh transgenic line in an embryo laying chamber with a grape juice agar plate and yeast paste. Flies were left for 2-3 days to mate and acclimate to the chamber. On the morning of collections, flies were first flipped onto fresh yeast paste while also anesthetizing with CO2 to encourage egg deposition and ensure accurate developmental timing. Two hours later, flies were flipped onto a fresh grape juice agar plate with minimal yeast paste for one hour and then removed. 14-15 hours later, embryos were sorted based on YFP fluorescence in the mouth hooks and processed as needed. All crosses and egg development were done at 25℃.

### Cuticle preparations and image processing

For cuticle preparations, non-YFP fluorescent embryos that were unhatched ∼30 hours after egg laying (AEL) were processed exactly as described in [53] with the following specifications: 20uL of 3:1 Lactic Acid/H2O was pipetted onto embryos, which were then incubated at 60℃ overnight. Darkfield images of five independently prepared cuticles per genotype were imaged on a Nikon E800 fluorescence microscope at 10x magnification. The ImageJ software was used to demarcate the boundaries of cuticle preparations by manually outlining and the aspect ratio was then automatically computed.

### Embryo viability assay

To assess embryonic viability, 150-200 non-YFP fluorescent embryos (∼15-16 hour AEL) were transferred to a fresh grape juice agar plate in groups of 10 to make counting easier. At 30 hours AEL, the percentage of hatched embryos from these 150-200 embryos was calculated. Importantly, this was repeated each day for no more than a week to avoid parental age-dependent effects on embryonic viability [54].

### Immunofluorescence and imaging

For immunofluorescence, non-YFP fluorescent embryos (∼15-16 hour AEL) with a FLAG-tagged Grh transgene were dechorionated in 11-15% hypochlorite. Embryos were then fixed and prepared for imaging. Briefly, embryos were fixed by rocking in ∼3.7% paraformaldehyde for 15 minutes at room temperature, the fixative was replaced with 100% methanol and embryos were shaken vigorously for 2 minutes. Embryos that had sunk to the bottom were washed with methanol and then stored in methanol at -80℃ until ready for processing.

For immunostaining, embryos were rehydrated with a methanol: PBST (1x PBS + 0.05% Triton-X) dilution series, washed in PBST for 30 minutes, blocked in 100uL Everyday Blocking Buffer (BioRad:12010020) for another 30 minutes, and then 1uL of a 1:100 dilution of an antibody raised against the FLAG epitope (abcam: ab205606) was added and embryos were rocked at 4℃ overnight. The next day, the antibody solution was removed and embryos were washed with PBST 3 times for 5 minutes each and then 2 times for 30 minutes each. Embryos were then incubated with 500uL of PBST mixed with 0.5uL of (555nm) fluorophore-conjugated secondary antibody (Invitrogen: A-21428) recognizing the FLAG primary antibody. Tubes were shielded from light and rocked for 2 hours at room temperature. Finally, embryos were washed with PBST 3 times for 5 minutes each and then 4 times for 30 minutes each, cleared and stained with 50% glycerol + DAPI (1ug/uL) for 1 hour, and then mounted with DABCO mounting media. Mounted embryos and single focal planes of the epidermis were imaged with a Leica SP5 confocal microscope. For the whole embryo image, Z-stacks and maximum intensity projections were produced using the Leica software, and images were then processed in ImageJ software (**Figure S8**).

### ATAC- and RNA-seq extractions and library preparations

For ATAC- and RNA-seq, 40-50 non-YFP fluorescent embryos (∼15-16 hour AEL) were dechorionated in 11-15% hypochlorite, snap-frozen in liquid nitrogen, and stored at -80℃. Chromatin for ATAC-seq and RNA for RNA-seq were extracted from 40-50 embryos per sample (2 biological replicates per genotype; only one transgenic replicate was used per genotype) using the protocol described in [51]. ChIP- and ATAC-seq libraries were prepared using the NEB Ultra II DNA Library Prep Kit according to the manufacturer’s instructions. RNA-seq libraries were prepared using the Illumina mRNA stranded kit according to the manufacturer’s instructions.

### Bioinformatic analyses

All reads were trimmed for adapters and quality checked using *trimgalore* [55]. Reads were then aligned to the *D. melanogaster* dm6 reference genome using *bowtie2-align* [56], sorted, and indexed using *samtools-sort* [57]. Duplicate reads were removed using *samtools-rmdup*. To identify regions enriched for ATAC-seq signal, we used the *macs2-callpeak* function [58] and then used the *BEDtools* merge function [59] to create a consensus peak set for each data type. Aligned reads that overlapped exonic regions for RNA-seq or consensus peaks for the ATAC-seq were counted for each sample using the *BEDtools* multicov function [59]. The resulting count matrices were used with *DEseq2* [60] to test for statistically significant differences (q < 0.05) in ATAC- and RNA-seq signals between samples. These data were visualized with *ggplot2* [61] and *deepTools* [62]. Lastly, for the fly-Grh direct targets, motif enrichments were identified using *icisTarget* [63] and gene ontology enrichments using the *GOslim* [39] molecular processes annotation database. See Github for all code used in analyses ( https://github.com/WittkoppLab/Ertl_et_al_Grh_rescue).

### Percent fly-Grh rescue

Percent rescue relative to the fly-Grh allele was calculated as follows:

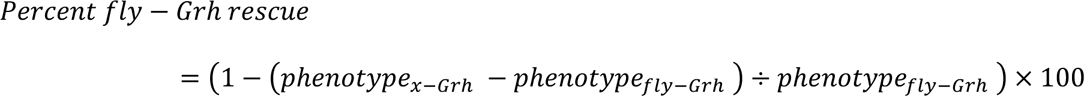

Where, *phenotype_x-Grh_* = the phenotypic measurement (accessibility or expression) for embryos expressing the experimental Grh transgene, and *phenotype_fly-Grh_* = the phenotypic measurement (accessibility or expression) for embryos expressing fly-Grh transgene.

## Supporting information

Other_Supplemental

## ACKNOWLEDGEMENTS

The authors thank members of the Wittkopp lab for feedback and research support, as well as Bill McGinnis for providing the *grh^IM^* line used.

## SUPPLEMENTARY INFORMATION

The supplementary information PDF file includes Supplementary Figures S1 - S8 and Supplementary Tables S1 - S4. Table S5 containing primers used is provided as an excel spreadsheet file. Plasmid maps for each Grh version transgenically inserted as well as the code used for all analyses (also available on GitHub https://github.com/WittkoppLab/Ertl_et_al_Grh_rescue) are provided in an additional folder titled: other_supplemental

**Figure S1.**
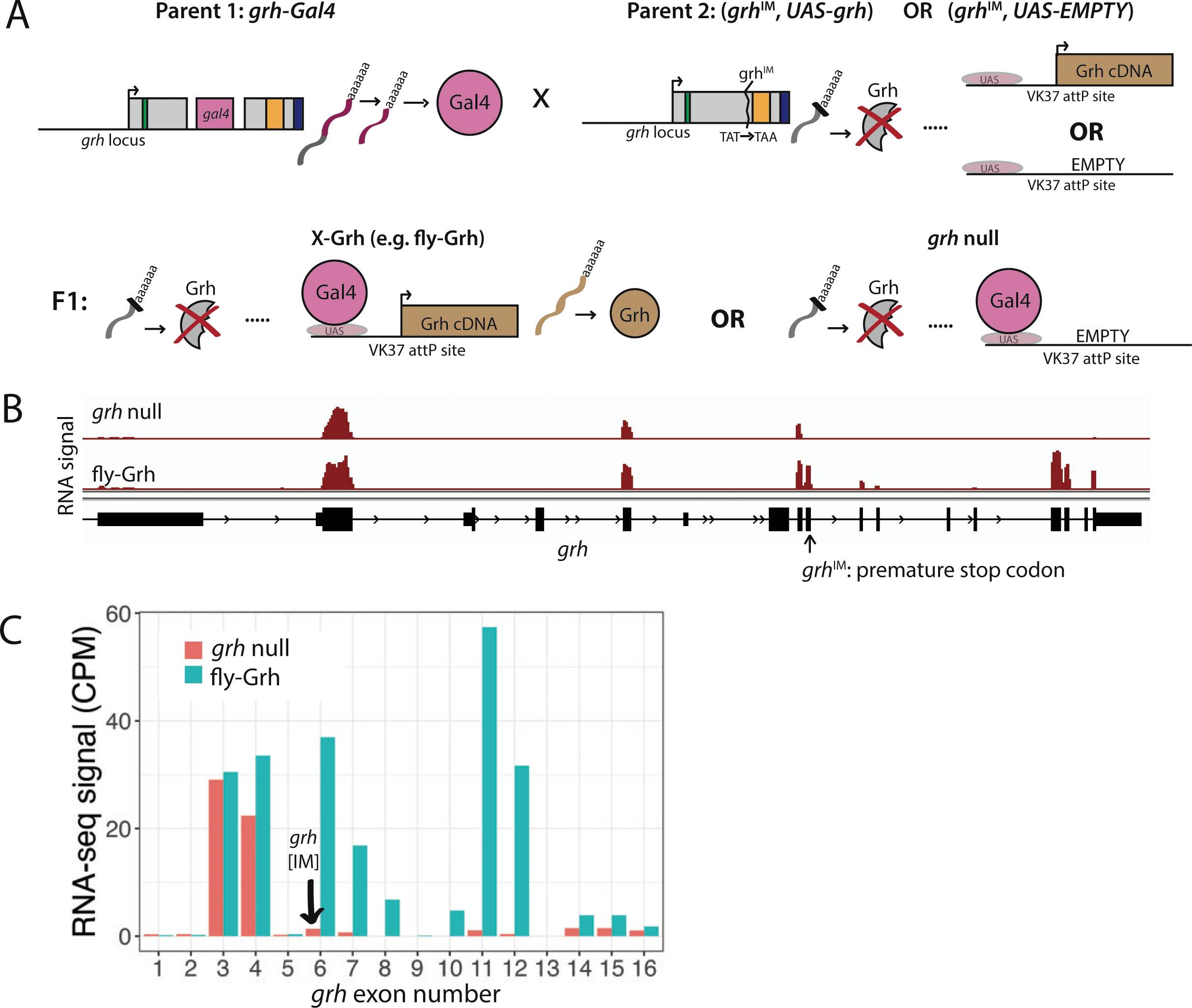
Experimental approach to test function of Grh transgenes in *grh* mutant embryos. **(A)** Schematic of the cross between ‘Parent 1’ (insertion of Gal4 coding sequence into the native *grh* locus to produce grh-Gal4, as described in Lee et al. 2018) and ‘Parent 2’ (UAS-transgene in a grh null background) to produce F1 experimental embryos that express any *grh* transgene (X-Grh) or an empty UAS vector (*grh* null). The Gal4 protein binds to the UAS to activate expression of Grh cDNA in the native *grh* expression profile. (**B**) Screenshot of genome browser of normalized RNA-seq data from *grh* null and fly-grh embryos. In contrast to fly-Grh embryos, *grh* null embryos show near-complete elimination of endogenous *grh* transcripts after the premature stop codon. (**C**) Quantification of (B) showing the total normalized RNA-seq read count per exon for fly-Grh and *grh* null embryos. The exon containing the premature stop codon is indicated.

**Figure S2.**
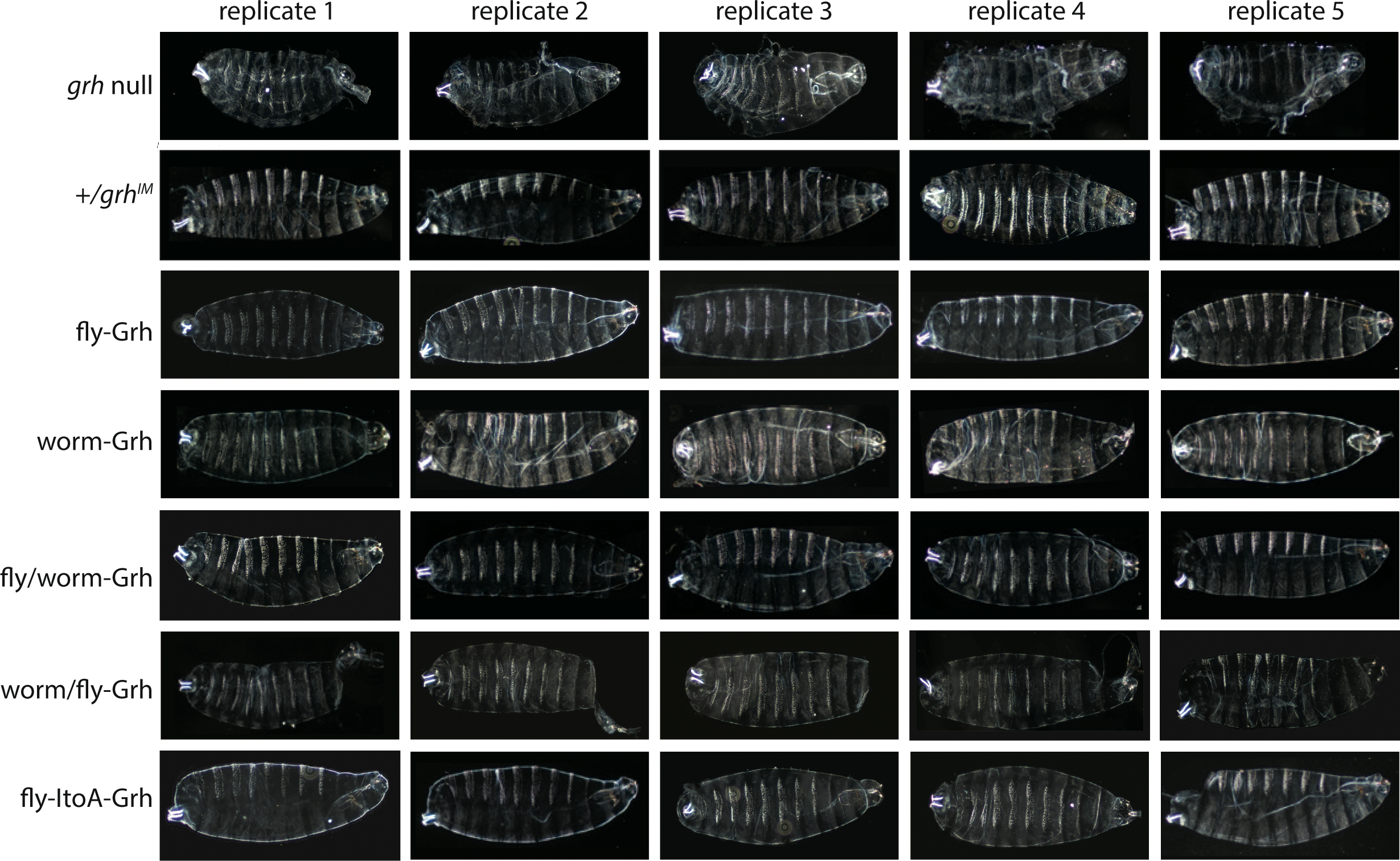
Cuticle preparations for all Grainy head transgenic lines. Five replicates of cuticle preparations for each of the Grh transgenic lines. Images were analyzed in ImageJ to create Figure 2F (see Methods).

**Figure S3.**
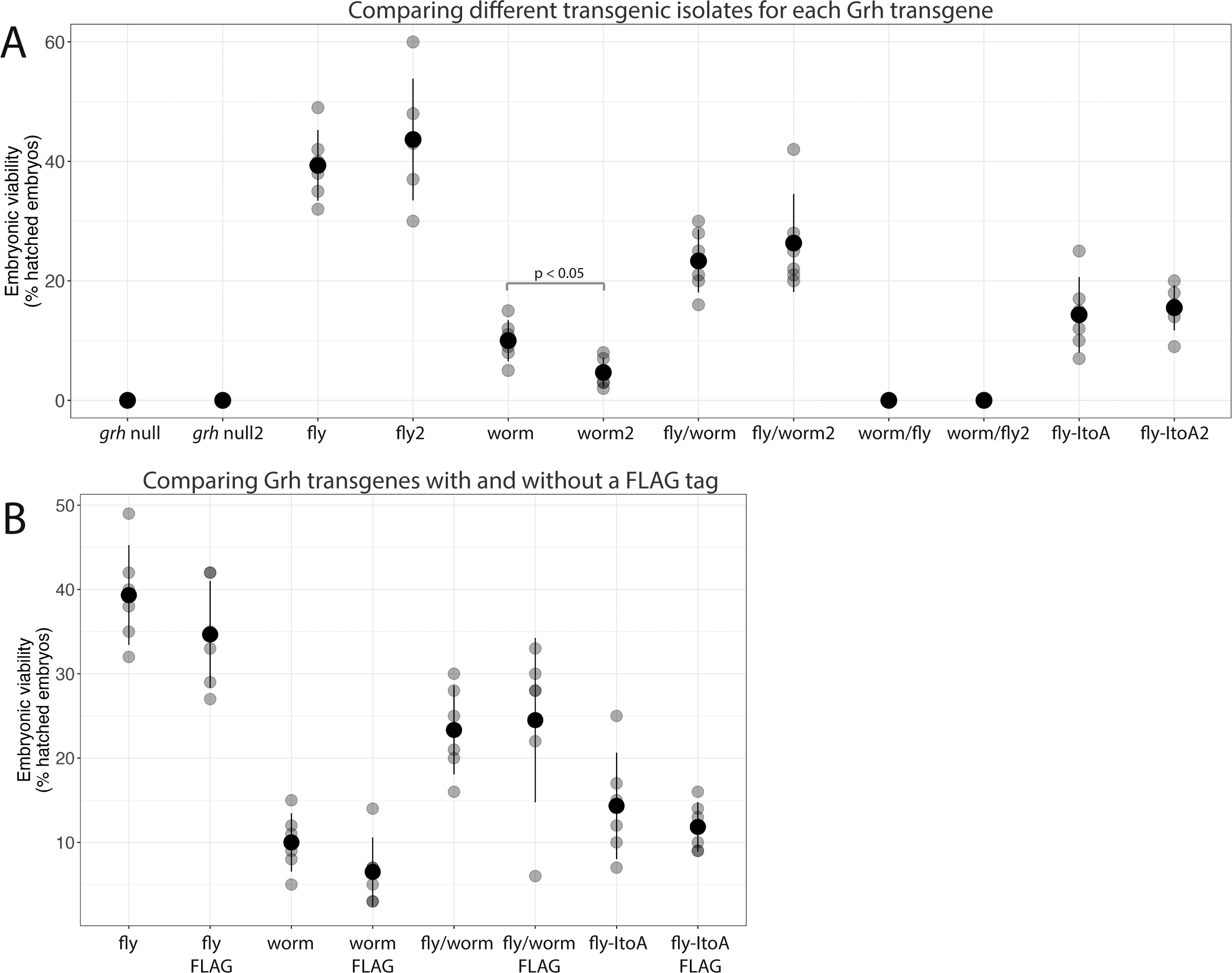
Embryonic viability for all Grh transgenic lines. Embryonic viability, measured as percent hatched of 150-200 embryos as in Figure 3, but comparing (A) two independent transgenic isolates and (B) transgenic lines with and without a FLAG tag on the (-terminus. All paired comparisons were not significantly different with the exception of the two isolates of the worm transgene: worm and worm2 (p < 0.05, Wilcoxon ranked sum test).

**Figure S4.**
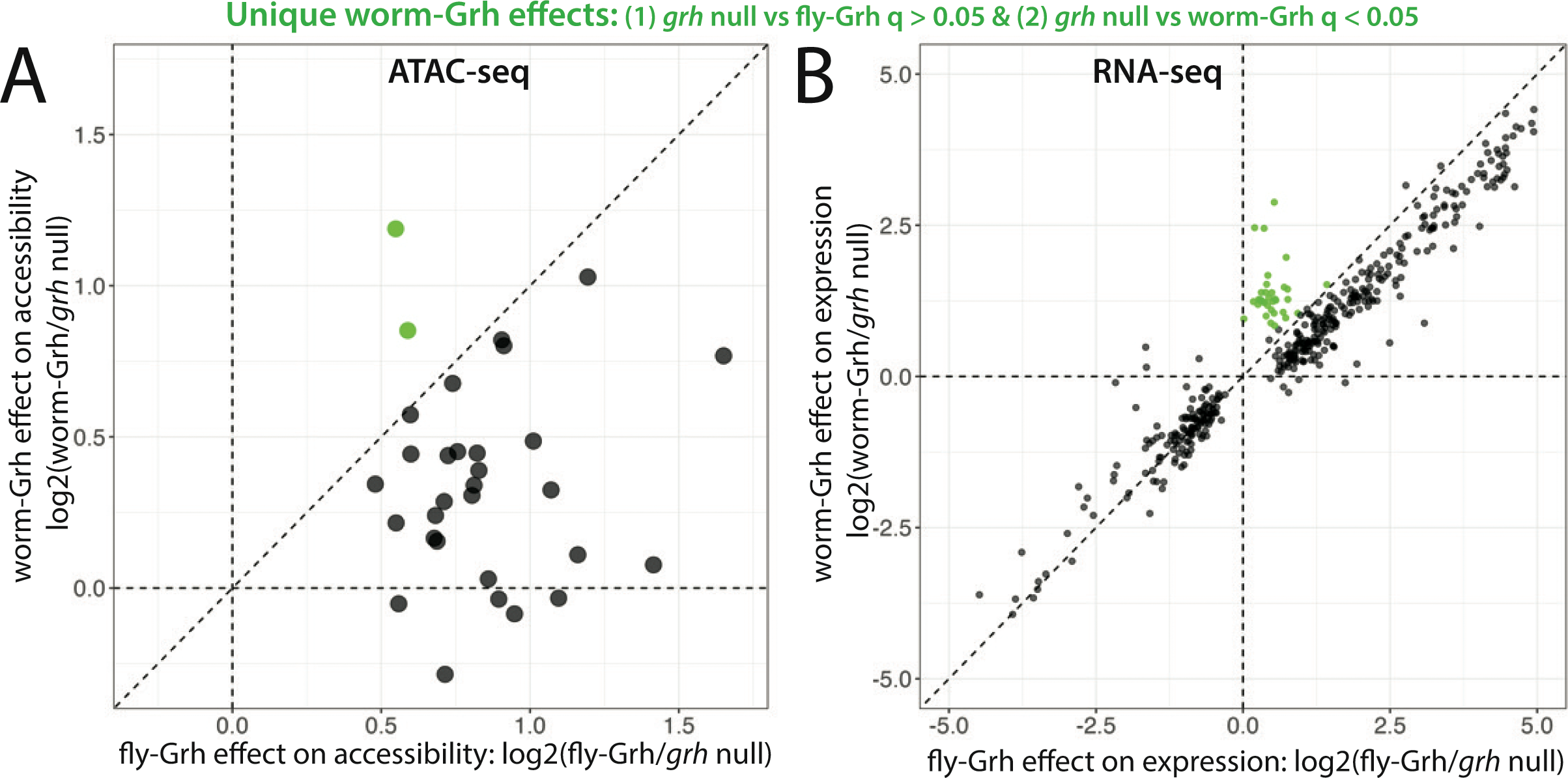
ATAC-seq processing and total accessibility comparison. **(A)** Flowchart showing worktlow for processing ATAC-seq data from reads to regions with statistically significant differential accessibility across transgenic lines. (**B**) Scatterplot contrasting the mean CPM (between two replicates) of accessibility between embryos expressing tly-Grh or worm-Grh for all accessible regions.

**Figure S5.**
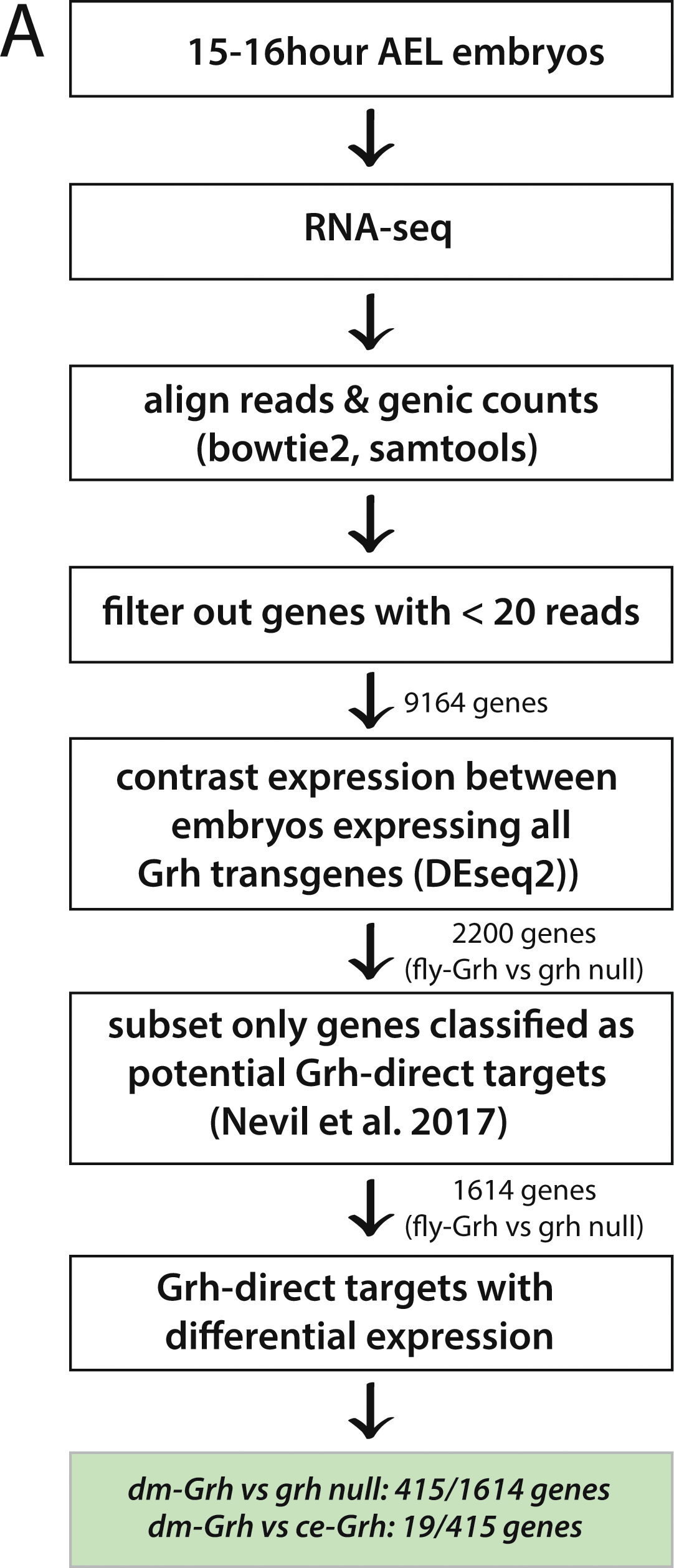
Unique worm-Grh effects on chromatin accessibility and gene expression. **(A)** Same scatterplot as shown in Figure 4E contrasting fly- and worm-Grh effects on chromatin accessibility, but also including regions for which q < 0.05 between grh null embryos and worm-Grh but not fly-Grh embryos (green). (**B**) Same scatterplot as shown in Figure 5C contrasting fly- and worm-Grh effects on gene expression, but also including regions for which q < 0.05 between grh null embryos and worm-Grh but not fly-Grh embryos (green).

**Figure S6.**
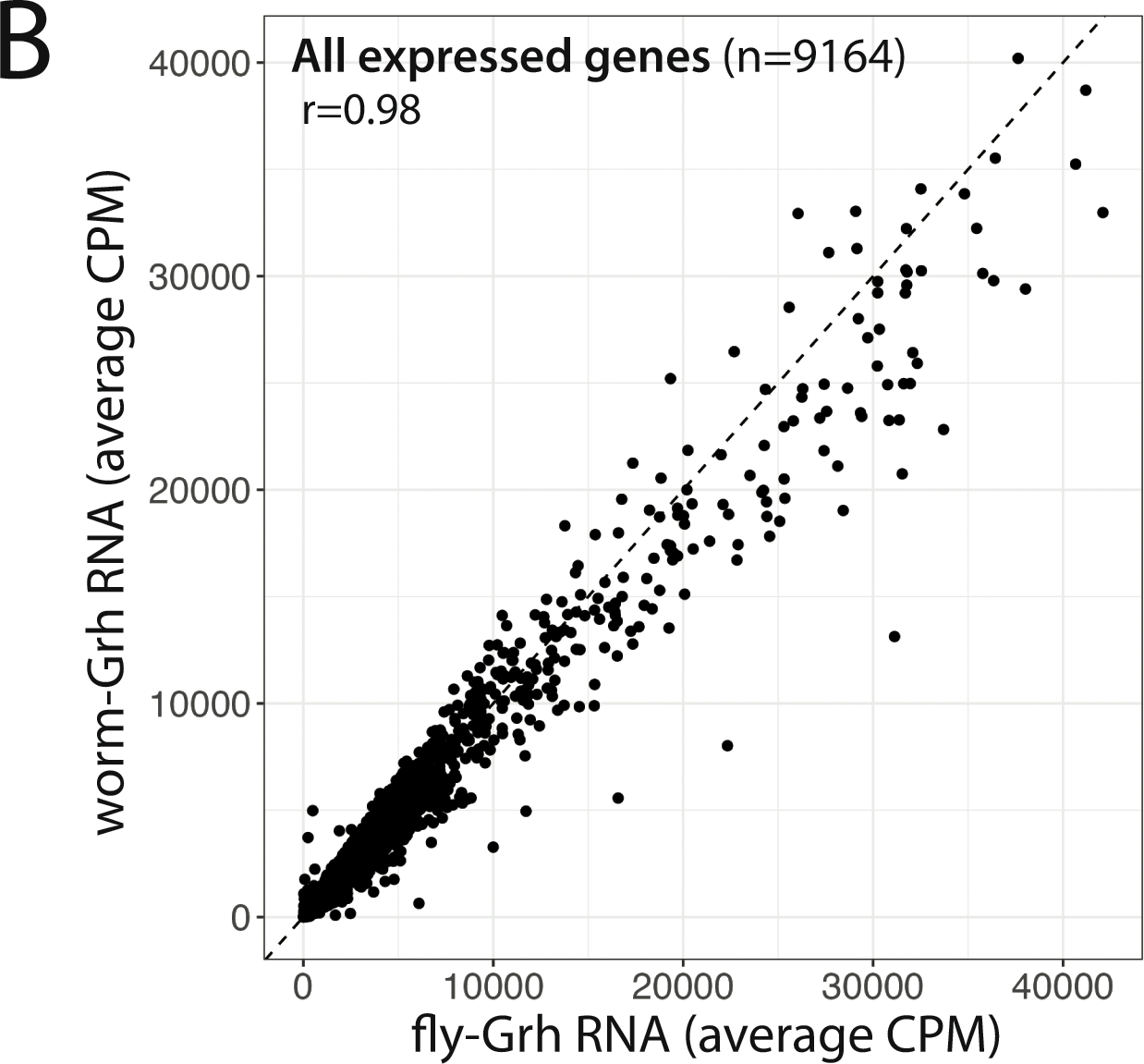
RNA-seq processing and total accessibility comparison. **(A)** Flowchart showing workflow for processing RNA-seq data from reads to regions with statistically significant differential gene expression across transgenic lines. (**B**) Scatterplot contrasting the mean CPM (between two replicates) of mRNA abundance of all expressed genes between embryos expressing fly-Grh or worm-Grh.

**Figure S7.**
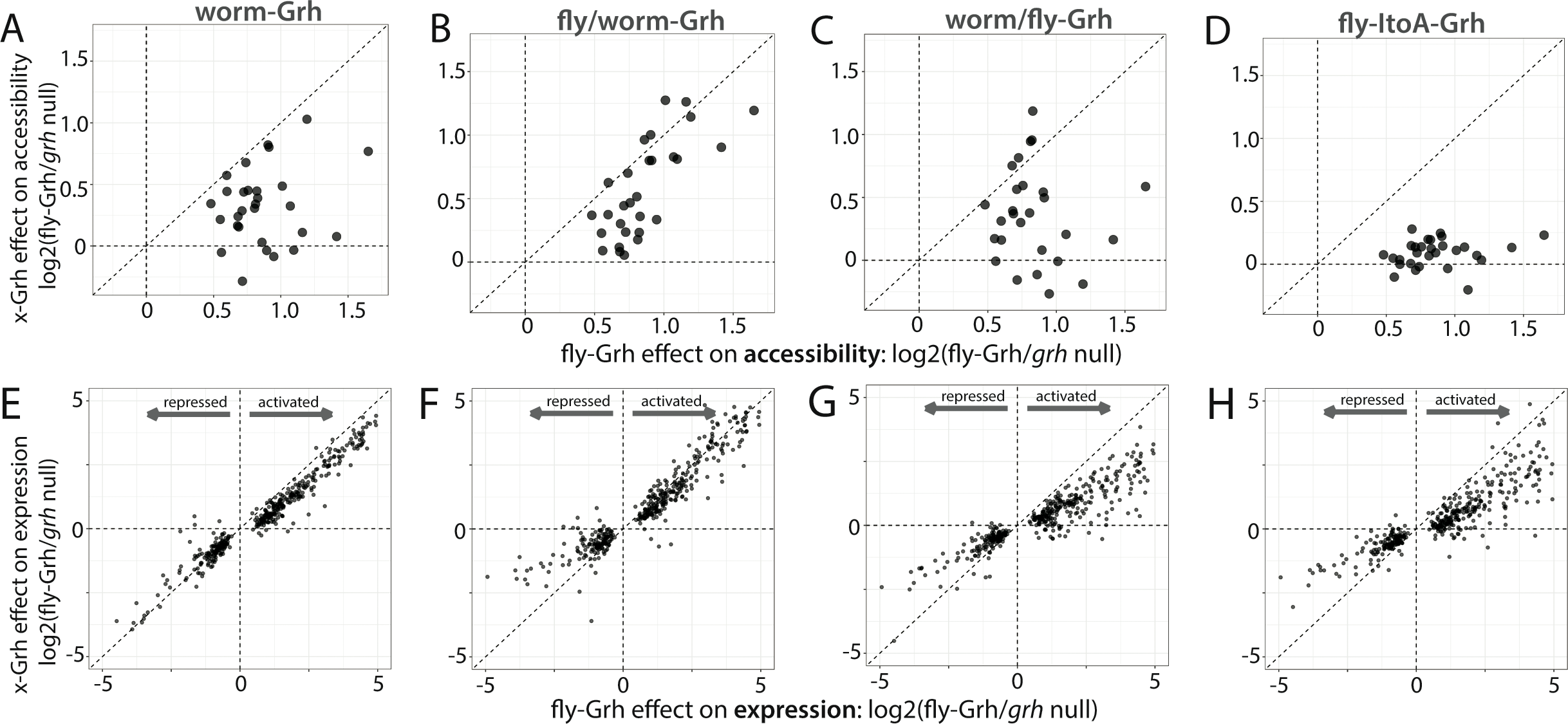
Chromatin accessibility and gene expression in grh null mutants expressing each of the Grainy head transgenes. (**A-D**} Scatterplots of fly-Grh effect on accessibility (log2(fly-Grh/grh nulll)l) (x-axisl) against that of all other Grainy head transgenes (specific Grh allele listed above each plotl). (**E-H**} Scatterplots of fly-Grh effect on expression (log2(fly-Grh/grh nulll)l) (x-axisl) against that of all other Grainy head transgenes (specific Grh allele listed above each plotl).

**Figure S8.**
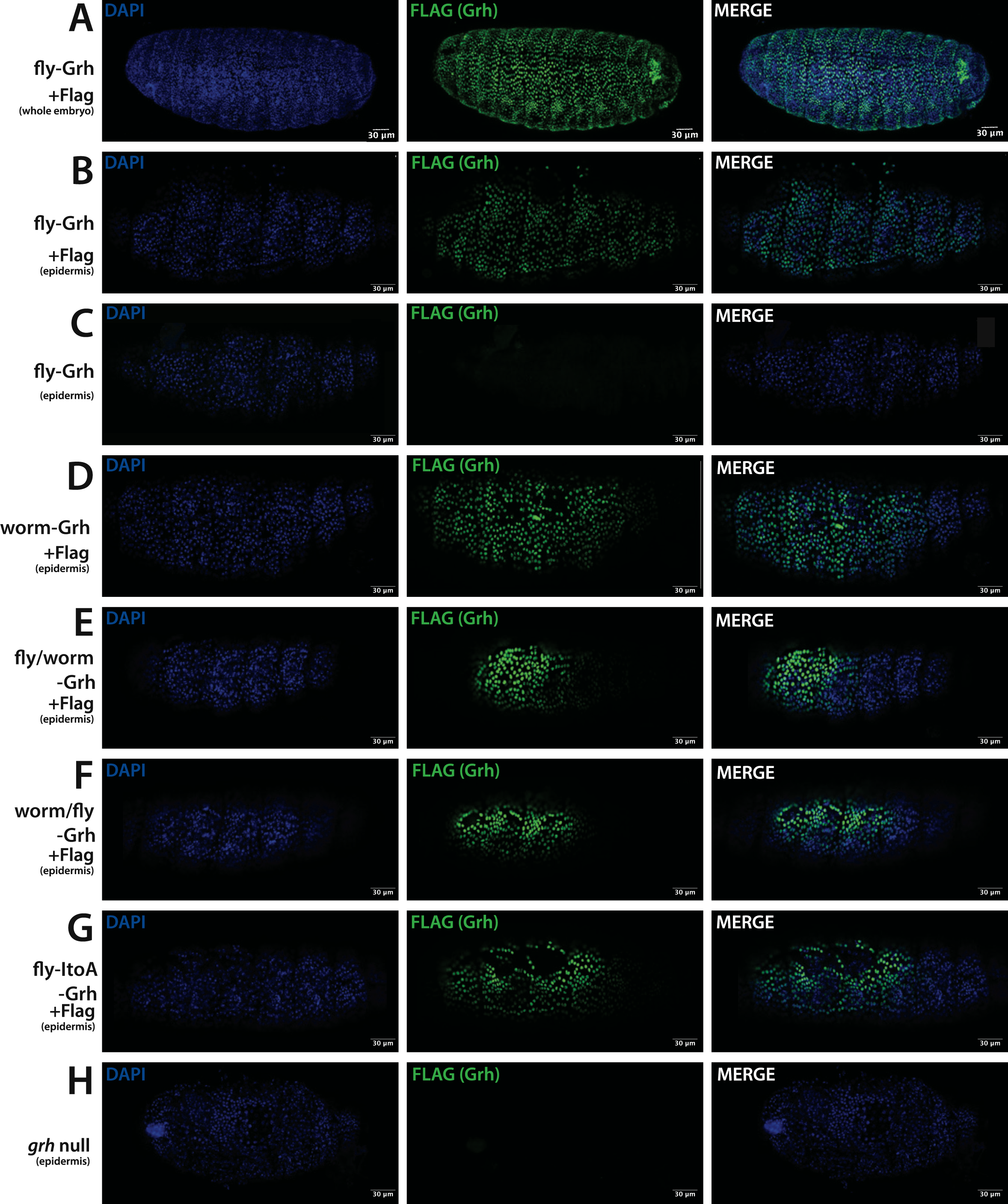
Grainy head transgenic proteins are translated in the embryonic epidermis. Confocal images of, from left to right, DAPI (DNA) in blue, FLAG (Grh) in green, and merged DAPI and FLAG signals for (A) a whole embryo expressing fly-Grh-Flag (whole embryo, Z-stack with maximum projection; see Methods) and epidermal cells (that is, a single focal plane of outer embryo) from embryos expressing (B) fly-Grh-Flag (epidermis), (C) fly-Grh (epidermis), (D) worm-Grh-Flag (epidermis), (E) fly/worm-Grh-Flag (epidermis), (F) worm/fly-Grh (epidermis), (G) fly-ItoA-Grh-Flag (epidermis), and (H) grh null embryos (epidermis).

**Table S1.** Summary of P values for aspect ratio. Values are from pairwise, post-hoc Tukey tests following ANOVA comparing the shape of cuticles from embryos expressing each allele of Grh analyzed. Shaded white = P > 0.05; orange = P < 0.05; yellow = P < 0.01.

| p value (FDR adj) | *grh* null | fly-Grh | worm-Grh | fly/worm-Grh | worm/fly-Grh | fly-ItoA-Grh |
| --- | --- | --- | --- | --- | --- | --- |
| *grh* null |  |  |  |  |  |  |
| fly-Grh | 0.000003 |  |  |  |  |  |
| worm-Grh | 0.000341 | 0.562541 |  |  |  |  |
| fly/worm-Grh | 0.000128 | 0.783305 | 0.999767 |  |  |  |
| worm/fly-Grh | 0.04963 | 0.012394 | 0.456372 | 0.263468 |  |  |
| fly-ItoA-Grh | 0.000223 | 0.661957 | 0.999998 | 0.9999917 | 0.3658018 |  |
| *grh*<sup>IM</sup>/+ | 0.000032 | 0.968394 | 0.97157 | 0.9999983 | 0.099303 | 0.9895581 |

**Table S2.** Summary of P values for aspect ratio. Values are from pairwise, post-hoc Tukey tests following ANOVA comparing the embryonic viability of embryos expressing each allele of Grh analyzed. Shaded white = P > 0.05; orange = P < 0.05; yellow = P < 0.01.

| p value (FDR adj) | *grh* null | fly-Grh | worm-Grh | fly/worm-Grh | worm/fly-Grh | fly-ItoA-Grh |
| --- | --- | --- | --- | --- | --- | --- |
| *grh* null |  |  |  |  |  |  |
| fly-Grh | 0 |  |  |  |  |  |
| worm-Grh | 0.3402459 | 0 |  |  |  |  |
| fly/worm-Grh | 0.0000004 | 0.002593 | 0.0328998 |  |  |  |
| worm/fly-Grh | 1 | 0 | 0.3402459 | 0.0000004 |  |  |
| fly-ItoA-Grh | 0.0134234 | 0 | 0.999277 | 0.5352427 | 0.0134234 |  |
| *grh*<sup>IM</sup>/+ | 0 | 0.00183 | 0 | 0 | 0 | 0 |

**Table S3.**
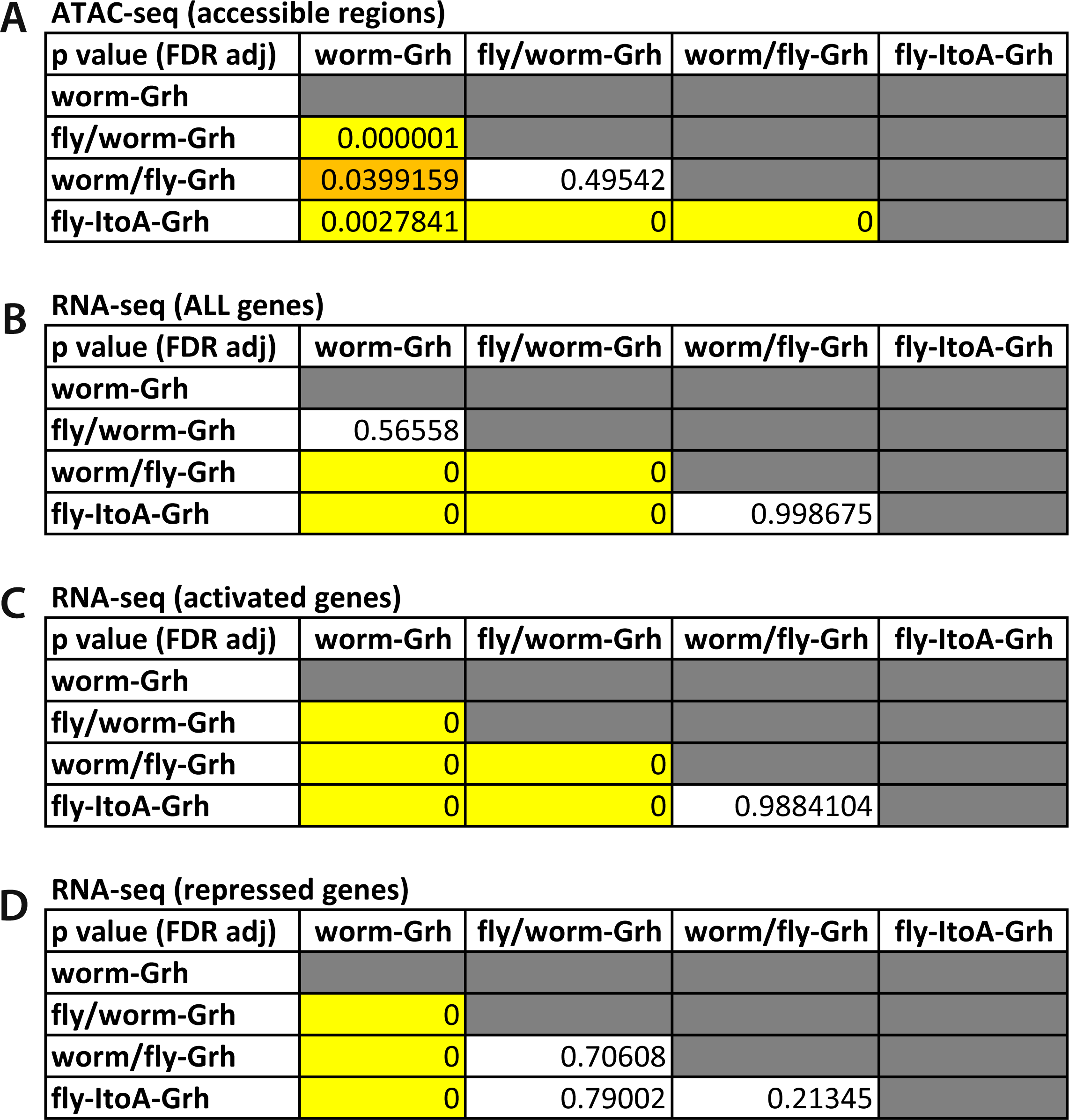
Summary of P values from functional genomics outputs. Values are from pairwise, post-hoc Tukey tests following ANOVA comparing (A) chromatin accessibility and (B-D) gene expression of (B) all, (C) activated, and (D) repressed Grh-regulated genes in embryos expressing each allele of Grh analyzed. Shaded white = P > 0.05; orange = P < 0.05; yellow = P < 0.01.

**A ATAC-seq (accessible regions)**
| p value (FDR adj) | worm-Grh | fly/worm-Grh | worm/fly-Grh | fly-ItoA-Grh |
| --- | --- | --- | --- | --- |
| worm-Grh |  |  |  |  |
| fly/worm-Grh | 0.000001 |  |  |  |
| worm/fly-Grh | 0.0399159 | 0.49542 |  |  |
| fly-ItoA-Grh | 0.0027841 | 0 | 0 |  |

**B RNA-seq (ALL genes)**
| p value (FDR adj) | worm-Grh | fly/worm-Grh | worm/fly-Grh | fly-ItoA-Grh |
| --- | --- | --- | --- | --- |
| worm-Grh |  |  |  |  |
| fly/worm-Grh | 0.56558 |  |  |  |
| worm/fly-Grh | 0 | 0 |  |  |
| fly-ItoA-Grh | 0 | 0 | 0.998675 |  |

**C RNA-seq (activated genes)**
| p value (FDR adj) | worm-Grh | fly/worm-Grh | worm/fly-Grh | fly-ItoA-Grh |
| --- | --- | --- | --- | --- |
| worm-Grh |  |  |  |  |
| fly/worm-Grh | 0 |  |  |  |
| worm/fly-Grh | 0 | 0 |  |  |
| fly-ItoA-Grh | 0 | 0 | 0.9884104 |  |

**Table S3. Summary of P values from functional genomics outputs.**
| p value (FDR adj) | worm-Grh | fly/worm-Grh | worm/fly-Grh | fly-ItoA-Grh |
| --- | --- | --- | --- | --- |
| worm-Grh |  |  |  |  |
| fly/worm-Grh | 0 |  |  |  |
| worm/fly-Grh | 0 | 0.70608 |  |  |
| fly-ItoA-Grh | 0 | 0.79002 | 0.21345 |  |

**Table S4.**
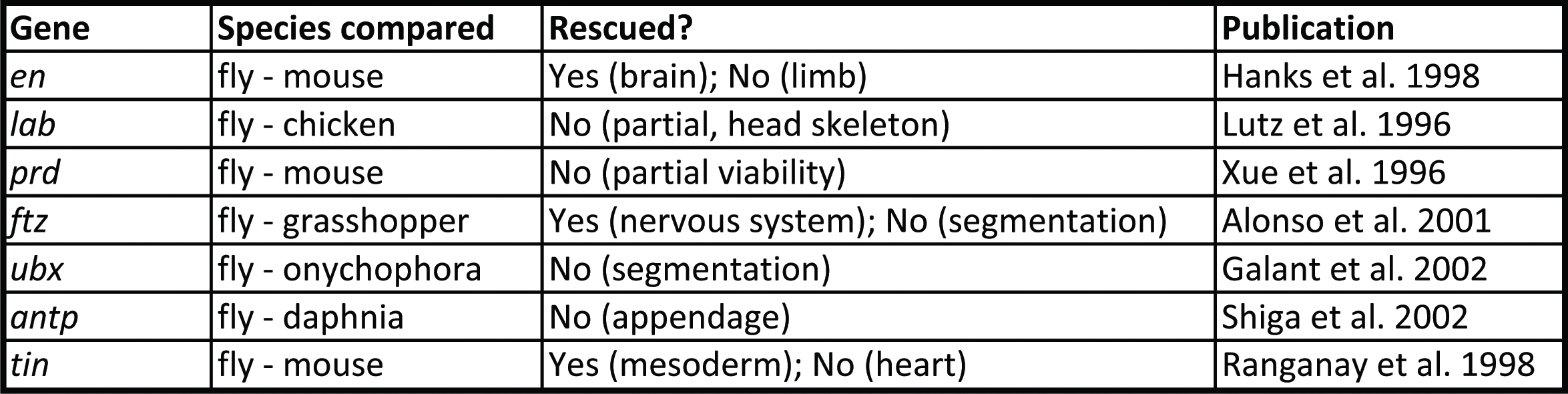
Summary of prior functional comparisons of divergent transcription factors.

