## Supplementary material for "Divergence of Grainy head affects chromatin accessibility, gene expression, and embryonic viability in *Drosophila melanogaster*": Other_Supplemental: code_for_analyses.R

```
## load libraries
library(plyr)
library(dplyr)
library(DESeq2)
library(ggplot2)
library(reshape2)
library(tidyr)

## load main plot theme
theme_main <- function() {
  theme_bw() +
  theme(
  #panel.grid.major = element_blank(),
  #panel.grid.minor = element_blank(),
  axis.text = element_text(size = 15),
  axis.title = element_text(size = 20),
  strip.text = element_text(size = 20),
  legend.text= element_text(size = 15),
  legend.title = element_text(size = 20),
  plot.title = element_text(size = 25, face = "bold")
)
}

#set working directory
setwd("/Users/henryertl/repos/Ertl_et_al_Grh_rescue")

#################################################################
############## Cuticle measurements -- Figure 2 #################
#################################################################

## read in df (from imageJ) and format
dfcut <- read.csv("/Users/henryertl/Downloads/cuticle_measurements.csv", header = T)
dfcut$grh <- c(rep("empty",5),rep("dm",5),rep("ce",5),rep("fw",5),rep("wf",5),rep("ia",5),rep("het",5))
dfcut[,1] <- NULL
dfcut$grh <- factor(dfcut$grh, levels=c("empty","dm","ce","het","fw","wf","ia"))

## plot main - Fig1
dfcut1 <- dfcut[dfcut$grh == "dm" | dfcut$grh == "ce" | dfcut$grh == "empty" | dfcut$grh == "het",]
cut_plot_1 <-  dfcut1 %>% na.omit() %>% ggplot(aes(x=grh, y=AR))+
geom_point(alpha = 0.3, size=3.5)+
stat_summary(fun.data=mean_sdl, fun.args = list(mult=1), geom="pointrange", color="black", size=1)+
theme_main()+
ylim(1.85,3.8)+
ylab("Aspect ratio)")+
xlab("")+
scale_x_discrete(labels=c('null','dm-Grh','ce-Grh', 'dm/ce-Grh', 'ce/dm-Grh', 'dm-Grh\n-ItoA','het'))+
theme(axis.text.x = element_text(size=15), axis.text.y = element_text(size=17), axis.title.y = element_text(size=17))
ggsave(cut_plot_1, file = "./plots/cuticle_AR_1.png", width = 4, height = 4)

## plot all - supplemental fig
cut_plot_2 <-  dfcut %>% na.omit() %>% ggplot(aes(x=grh, y=AR))+
geom_point(alpha = 0.3, size=5)+
stat_summary(fun.data=mean_sdl, fun.args = list(mult=1), geom="pointrange", color="black", size=1.5)+
theme_main()+
ylab("Aspect ratio)")+
xlab("")+
scale_x_discrete(labels=c('null','dm-Grh','ce-Grh', 'dm/ce-Grh', 'ce/dm-Grh', 'dm-Grh\n-ItoA','het'))+
theme(axis.text.x = element_text(size=15), axis.text.y = element_text(size=17), axis.title.y = element_text(size=17))
ggsave(cut_plot_2, file = "./plots/cuticle_AR_2.png")

# run ANOVA and Tukey to get stats for supplemental P val table
anova_cut <- aov(AR ~ grh, data = dfcut)
TukeyHSD(anova_cut, conf.level=.95)


#################################################################
############### Embryonic Viability -- Figure 3 #################
#################################################################

## read in df and format
hatch <- read.delim("./Counts/hatching_data.txt", header = T)
hatch_form <- melt(hatch)
hatch_form <- hatch_form[hatch_form$variable != "n" & hatch_form$variable != "mm" & hatch_form$variable != "n2" & hatch_form$variable != "mm2",]
hatch_form$variable <- factor(hatch_form$variable, levels=c("empty","empty2","dm","dm2","dmf1","dm2_bal_sib","ce","ce2","cef1","ce2_bal_sib","empty1_bal","fw","fw2","fwf1","wf","wf2","ia","ia2","iaf1"))

## plot main - Fig3
hatch_main1 <- hatch_form[hatch_form$variable == "empty" | hatch_form$variable == "dm" | hatch_form$variable == "ce" | hatch_form$variable == "empty1_bal",]
hatch_plot_main1 <-  hatch_main1 %>% na.omit() %>% ggplot(aes(x=variable, y=value))+
geom_point(alpha = 0.3, size=3.5)+
stat_summary(fun.data=mean_sdl, fun.args = list(mult=1), geom="pointrange", color="black", size=1)+
theme_main()+
ylim(0,80)+
ylab("Embryonic viability\n(% hatched embryos)")+
xlab("")+
scale_x_discrete(labels=c('grh null','dm-Grh','ce-Grh',"grh null\nhet."))+
theme(axis.text.x = element_text(size=15), axis.text.y = element_text(size=17), axis.title.y = element_text(size=17))
ggsave(hatch_plot_main1, file = "./plots/hatching_freq_main1.png", width = 4, height = 4)

## plot main - Fig6
hatch_main2 <- hatch_form[hatch_form$variable == "empty" | hatch_form$variable == "dm" | hatch_form$variable == "ce" | hatch_form$variable == "fw"| hatch_form$variable == "wf" | hatch_form$variable == "ia",]
hatch_plot_main2 <-  hatch_main2 %>% na.omit() %>% ggplot(aes(x=variable, y=value))+
geom_point(alpha = 0.3, size=5)+
stat_summary(fun.data=mean_sdl, fun.args = list(mult=1), geom="pointrange", color="black", size=1.5)+
theme_main()+
ylab("Embryonic viability\n(% hatched embryos)")+
xlab("")+
scale_x_discrete(labels=c('grh null','dm-Grh','ce-Grh', 'dm/ce-Grh', 'ce/dm-Grh', 'dm-Grh\n-ItoA'))+
theme(axis.text.x = element_text(size=15), axis.text.y = element_text(size=17), axis.title.y = element_text(size=17))
ggsave(hatch_plot_main2, file = "./plots/hatching_freq_main2.png", width = 8, height = 6)

## plot supp figure -- transgenic replicates
hatch_supp_12 <- hatch_form[hatch_form$variable != "dmf1" & hatch_form$variable != "cef1" & hatch_form$variable != "fwf1" & hatch_form$variable != "iaf1" & hatch_form$variable != "empty1_bal" & hatch_form$variable != "dm2_bal_sib" & hatch_form$variable != "ce2_bal_sib",]
hatch_plot_supp12 <-  hatch_supp_12 %>% na.omit() %>% ggplot(aes(x=variable, y=value))+
geom_point(alpha = 0.3, size=5)+
stat_summary(fun.data=mean_sdl, fun.args = list(mult=1), geom="pointrange", color="black", size=1.5)+
theme_main()+
ylab("Embryonic viability\n(% hatched embryos)")+
xlab("")+
scale_x_discrete(labels=c('grh null','grh null2','dm-Grh','dm-Grh2','ce-Grh', 'ce-Grh2','dm/ce-Grh', 'dm/ce-Grh2','ce/dm-Grh','ce/dm-Grh2', 'dm-Grh\n-ItoA','dm-Grh\n-ItoA2'))+
theme(axis.text.x = element_text(size=15), axis.text.y = element_text(size=17), axis.title.y = element_text(size=17))
ggsave(hatch_plot_supp12, file = "./plots/hatching_freq_supp12.png", width = 16)

## plot supp figure -- FLAG v no FLAG
hatch_supp_flag <- hatch_form[hatch_form$variable == "dm" | hatch_form$variable == "ce" | hatch_form$variable == "fw" | hatch_form$variable == "ia" | hatch_form$variable == "dmf1" | hatch_form$variable == "cef1" | hatch_form$variable == "fwf1"| hatch_form$variable == "iaf1",]
hatch_plot_flag <-  hatch_supp_flag %>% na.omit() %>% ggplot(aes(x=variable, y=value))+
geom_point(alpha = 0.3, size=5)+
stat_summary(fun.data=mean_sdl, fun.args = list(mult=1), geom="pointrange", color="black", size=1.5)+
theme_main()+
ylab("Embryonic viability\n(% hatched embryos)")+
xlab("")+
scale_x_discrete(labels=c('dm-Grh','dm-Grh\nFLAG','ce-Grh', 'ce-Grh\nFLAG','dm/ce-Grh', 'dm/ce-Grh\nFLAG', 'dm-Grh\n-ItoA','dm-Grh\n-ItoA\nFLAG'))+
theme(axis.text.x = element_text(size=15), axis.text.y = element_text(size=17), axis.title.y = element_text(size=17))
ggsave(hatch_plot_flag, file = "./plots/hatching_freq_flag.png", width = 10)

# run ANOVA and Tukey to get stats for supplemental P val table
anova_embryo <- aov(value ~ variable, data = hatch_form)
summary(anova_embryo )
TukeyHSD(anova_embryo , conf.level=.95)


#################################################################
######## Chromatin accessibility/ATAC-seq -- Figure 4 ###########
#################################################################

## read in dataframe and clean up
df <- read.delim("./counts/grh_atac_counts.txt", header = F)
df$locus <- paste(df$V1, df$V2, df$V3, sep = "_")
df[,c(1:3)] <- NULL
rawCounts_atac <- NULL
rawCounts_atac <- df
colnames(rawCounts_atac) <- c("dmF1","dmF2","ceF1","ceF2","mF1","mF2","empty1","empty2","IAF1","IAF2","NF1","NF2","FWF1","FWF2","WFF1","WFF2","locus")
rownames(rawCounts_atac) <- rawCounts_atac$locus
rawCounts_atac$locus <- NULL

## read in file key
sampleData <- read.delim("./other_files/name_order.txt")
sampleData$genotype <- sampleData$run
sampleData$run <- rownames(sampleData)

# run DEseq2
deseq2Data_atac <- DESeqDataSetFromMatrix(countData=rawCounts_atac, colData=sampleData, design= ~ genotype)
deseq2Data_atac <- DESeq(deseq2Data_atac)

# get DEseq2 values from all datasets
wt_dm <- results(deseq2Data_atac, contrast=c("genotype", "dm", "em")) %>% as.data.frame() %>% na.omit()
wt_dm$gene <- row.names(wt_dm)
wt_dm_sub <- wt_dm
colnames(wt_dm_sub) <- paste(colnames(wt_dm_sub), "dm", sep = "_")
wt_dm_sub$gene <- wt_dm_sub$gene_dm

wt_ce <- results(deseq2Data_atac, contrast=c("genotype", "ce", "em")) %>% as.data.frame() %>% na.omit()
wt_ce$gene <- row.names(wt_ce)
wt_ce_sub <- wt_ce
colnames(wt_ce_sub) <- paste(colnames(wt_ce_sub), "ce", sep = "_")
wt_ce_sub$gene <- wt_ce_sub$gene_ce

wt_fw <- results(deseq2Data_atac, contrast=c("genotype", "fw", "em")) %>% as.data.frame() %>% na.omit()
wt_fw$gene <- row.names(wt_fw)
wt_fw_sub <- wt_fw
colnames(wt_fw_sub) <- paste(colnames(wt_fw_sub), "fw", sep = "_")
wt_fw_sub$gene <- wt_fw_sub$gene_fw

wt_ia <- results(deseq2Data_atac, contrast=c("genotype", "ia", "em")) %>% as.data.frame() %>% na.omit()
wt_ia$gene <- row.names(wt_ia)
wt_ia_sub <- wt_ia
colnames(wt_ia_sub) <- paste(colnames(wt_ia_sub), "ia", sep = "_")
wt_ia_sub$gene <- wt_ia_sub$gene_ia

wt_wf <- results(deseq2Data_atac, contrast=c("genotype", "wf", "em")) %>% as.data.frame() %>% na.omit()
wt_wf$gene <- row.names(wt_wf)
wt_wf_sub <- wt_wf
colnames(wt_wf_sub) <- paste(colnames(wt_wf_sub), "wf", sep = "_")
wt_wf_sub$gene <- wt_wf_sub$gene_wf

dm_ce <- results(deseq2Data_atac, contrast=c("genotype", "dm", "ce")) %>% as.data.frame() %>% na.omit()
dm_ce$gene <- row.names(dm_ce)
dm_ce_sub <- dm_ce
colnames(dm_ce_sub) <- paste(colnames(dm_ce_sub), "dm_ce", sep = "_")
dm_ce_sub$gene <- dm_ce_sub$gene_dm_ce

# combine all datasets
all_atac <- join_all(list(wt_dm_sub, wt_ce_sub, wt_fw_sub, wt_ia_sub, wt_wf_sub, dm_ce_sub), by = 'gene', type='full') %>% na.omit() %>% unique()

# rename gene column
all_atac$locus <- all_atac$gene_wf
all_atac <- all_atac %>% separate(locus, c('chr', 'start','end'))
atac_small <- all_atac %>% select(chr, start, end, gene, log2FoldChange_dm, log2FoldChange_ce, log2FoldChange_fw, log2FoldChange_wf, log2FoldChange_ia, log2FoldChange_dm_ce, padj_dm, padj_ce, padj_fw, padj_wf, padj_ia, padj_dm_ce) %>% na.omit()
atac_small <- atac_small[atac_small$chr != "chrUn",]
atac_small <- atac_small[atac_small$end != "random",]

# assign as overlapping with grh chip peak or not
## read in dataset from Nevil et al 2017
chip <- read.delim(file = "./counts/grh_atac_counts_chipsub.txt", header = F)

## intersect and assign as yes or no overlapping
atac_grh <- bt.intersect(atac_small, chip, wao = TRUE)
atac_grh$V37 <- "NO"
for (i in 1:nrow(atac_grh)) {
if (atac_grh$V36[i] != 0){
	atac_grh$V37[i] <- "YES"
}
}
atac_grh <- atac_grh[,-c(17:36)]
colnames(atac_grh) <- c("chr", "start", "end", "gene", "log2FoldChange_dm", "log2FoldChange_ce", "log2FoldChange_fw", "log2FoldChange_wf", "log2FoldChange_ia", "log2FoldChange_dm_ce", "padj_dm", "padj_ce", "padj_fw", "padj_wf", "padj_ia", "padj_dm_ce","grh_overlap")

## write and read tables to clean up
write.table(atac_grh , file = "./counts/atac_grh_DEseq.txt", row.names = F, quote = F, sep = "\t")
atac_grh <- read.delim("./counts/atac_grh_DEseq.txt", header = T)
atac_grh_only <- atac_grh[atac_grh$grh_overlap == "YES" & atac_grh$padj_dm < 0.05,]
atac_grh_only_bed <- atac_grh_only[,c(1:3)]
write.table(atac_grh_only_bed , file = "./counts/atac_grh_DEseq_onlyGrh.txt", row.names = F, quote = F, sep = "\t")

# Volcano plot
atac_volc <- ggplot(atac_grh, aes(x=log2FoldChange_dm,y=-log10(padj_dm),color=grh_overlap))+
geom_point(alpha=0.6)+
xlim(-2,2)+
theme_main()+
xlab("log2(dm-Grh/grh null)")+
ylab("-log10(qvalue)")+
geom_hline(yintercept = -log10(0.05), linetype = "dashed", color = "orange") +
guides(colour = "none")
ggsave(atac_volc, file = "./plots/atac_volcano.png", width = 3.5, height= 3)

# Rescue plot
atac_rescue_ce <- atac_grh[atac_grh$padj_dm < 0.05 & atac_grh$grh_overlap == "YES",] %>%
ggplot(aes(x=log2FoldChange_dm, y=log2FoldChange_ce))+
geom_point(alpha=0.7, size=4.5)+
ylim(-0.3,1.7)+
xlim(-0.3,1.7)+
geom_hline(yintercept = 0, linetype = "dashed") +
geom_vline(xintercept = 0, linetype = "dashed") +
geom_abline(intercept = 0, slope = 1, linetype = "dashed") + theme_main() + geom_vline(xintercept = 0, linetype = "dashed")+
ylab("dm/ce-Grh effect on accessibility\nlog2(ce-Grh/grh null)")+
xlab("dm-Grh effect on accessibility\nlog2(dm-Grh/grh null)")
ggsave(atac_rescue_ce, file = "./plots/ce_rescue_ATAC_small_ce.png", height = 5.5, width = 6)

# load libraries for functional analyses to get pie chart
library(ChIPseeker)
library(TxDb.Dmelanogaster.UCSC.dm6.ensGene)
library(clusterProfiler)
library(AnnotationDbi)
# annotate and plot
edg <- TxDb.Dmelanogaster.UCSC.dm6.ensGene
peakAnno.edb <- annotatePeak(peaks, tssRegion=c(-3000, 3000),
                             TxDb=edb, annoDb="org.Dm.eg.db")


#################################################################
############# Gene expression/RNA-seq -- Figure 5 ###############
#################################################################

## read in df
df <- read.delim("./counts/grh_rna_counts.txt", header = F)
# compute genic counts from exonic
df_genic_counts <- ddply(df,.(V4),summarize,V5=sum(V5),V6=sum(V6),V7=sum(V7),V8=sum(V8),V9=sum(V9),V10=sum(V10),
V11=sum(V11),V12=sum(V12),V13=sum(V13),V14=sum(V14),V15=sum(V15),V16=sum(V16),
V17=sum(V17),V18=sum(V18),V19=sum(V19),V20=sum(V20))

write.table(df_genic_counts, file = "./counts/grh_rna_genic_counts.txt", sep = "\t", quote = F, row.names = F)

df_genic_counts <- read.delim("./counts/grh_rna_genic_counts.txt", header = T)
df_genic_counts <- df_genic_counts[df_genic_counts$V5 < 50000 & df_genic_counts$V5 > 20,]

# prepare files for DEseq2
rawCounts_RNA <- NULL
rawCounts_RNA <- df_genic_counts
colnames(rawCounts_RNA) <- c("gene","dmF1","dmF2","ceF1","ceF2","mF1","mF2","empty1","empty2","IAF1","IAF2","NF1","NF2","FWF1","FWF2","WFF1","WFF2")
rownames(rawCounts_RNA) <- rawCounts_RNA$gene
rawCounts_RNA$gene <- NULL

# read in sample key
sampleData <- read.delim("./other_files/name_order.txt")
sampleData$genotype <- sampleData$run
sampleData$run <- rownames(sampleData)

# run DEseq2
deseq2Data_RNA <- DESeqDataSetFromMatrix(countData=rawCounts_RNA, colData=sampleData, design= ~ genotype)
deseq2Data_RNA <- DESeq(deseq2Data_RNA)

# get DEseq2 values from all datasets vs empty
wt_dm <- results(deseq2Data_RNA, contrast=c("genotype", "dm", "em")) %>% as.data.frame() %>% na.omit()
wt_dm$gene <- row.names(wt_dm)
wt_dm_sub <- wt_dm
colnames(wt_dm_sub) <- paste(colnames(wt_dm_sub), "dm", sep = "_")
wt_dm_sub$gene <- wt_dm_sub$gene_dm

wt_ce <- results(deseq2Data_RNA, contrast=c("genotype", "ce", "em")) %>% as.data.frame() %>% na.omit()
wt_ce$gene <- row.names(wt_ce)
wt_ce_sub <- wt_ce
colnames(wt_ce_sub) <- paste(colnames(wt_ce_sub), "ce", sep = "_")
wt_ce_sub$gene <- wt_ce_sub$gene_ce

wt_fw <- results(deseq2Data_RNA, contrast=c("genotype", "fw", "em")) %>% as.data.frame() %>% na.omit()
wt_fw$gene <- row.names(wt_fw)
wt_fw_sub <- wt_fw
colnames(wt_fw_sub) <- paste(colnames(wt_fw_sub), "fw", sep = "_")
wt_fw_sub$gene <- wt_fw_sub$gene_fw

wt_ia <- results(deseq2Data_RNA, contrast=c("genotype", "ia", "em")) %>% as.data.frame() %>% na.omit()
wt_ia$gene <- row.names(wt_ia)
wt_ia_sub <- wt_ia
colnames(wt_ia_sub) <- paste(colnames(wt_ia_sub), "ia", sep = "_")
wt_ia_sub$gene <- wt_ia_sub$gene_ia

wt_wf <- results(deseq2Data_RNA, contrast=c("genotype", "wf", "em")) %>% as.data.frame() %>% na.omit()
wt_wf$gene <- row.names(wt_wf)
wt_wf_sub <- wt_wf
colnames(wt_wf_sub) <- paste(colnames(wt_wf_sub), "wf", sep = "_")
wt_wf_sub$gene <- wt_wf_sub$gene_wf

dm_ce <- results(deseq2Data_RNA, contrast=c("genotype", "dm", "ce")) %>% as.data.frame() %>% na.omit()
dm_ce$gene <- row.names(dm_ce)
dm_ce_sub <- dm_ce
colnames(dm_ce_sub) <- paste(colnames(dm_ce_sub), "dm_ce", sep = "_")
dm_ce_sub$gene <- dm_ce_sub$gene_dm_ce

# combine all datasets
all_rna <- join_all(list(wt_dm_sub, wt_ce_sub, wt_fw_sub, wt_ia_sub, wt_wf_sub, dm_ce_sub), by = 'gene', type='full') %>% na.omit() %>% unique()
# rename gene column
all_rna$gene <- all_rna$gene_wf

# subset only direct targets from Nevil et al 2017 (also need to convert IDs)
key <- read.delim("./other_files/Dmel_geneID_conversion_table.txt", header = F)
colnames(key) <- c("name", "spp", "fb", "fb2", "gene")
key <- key[,c(1,5)]
harrison <- read.delim("./other_files/grh_harrison_diff_express.txt", header = F)
colnames(harrison) <- "name"
harrison_final <- left_join(harrison, key, by = "name") %>% na.omit()

all_rna_final <- join_all(list(all_rna, harrison_final), by = 'gene', type = 'full')
all_rna_final$name[is.na(all_rna_final$name)] <- 0
all_rna_final$direct_target <- "NO"
for (i in 1:nrow(all_rna_final)) {
if (all_rna_final$name[i] != 0){
	all_rna_final$direct_target[i] <- "YES"
}
}

# clean up df and get coords
exon_coord <- read.delim("./other_files/dm6_gene_coords_first_exon.bed", header = F)
colnames(exon_coord) <- c("chr", "start", "end", "gene")
all_rna_final <- left_join(all_rna_final, exon_coord, by = "gene")
rna_small <- all_rna_final %>% select(chr, start, end, gene, log2FoldChange_dm, log2FoldChange_ce, log2FoldChange_fw, log2FoldChange_wf, log2FoldChange_ia, log2FoldChange_dm_ce, padj_dm, padj_ce, padj_fw, padj_wf, padj_ia, padj_dm_ce, direct_target) %>% na.omit()
rna_small_direct <- rna_small[rna_small$direct_target == "YES",]
write.table(rna_small , file = "./counts/rna_all_DEseq.txt", row.names = F, quote = F, sep = "\t")
write.table(rna_small_direct , file = "./counts/rna_all_DEseq_direct.txt", row.names = F, quote = F, sep = "\t")
rna_small_direct <- read.delim("./counts/rna_all_DEseq_direct.txt", header = T)

# Volcano plot
rna_volc <- ggplot(rna_small_direct, aes(x=log2FoldChange_dm,y=-log10(padj_dm)))+
geom_point(alpha=0.6)+
xlim(-5,5)+
theme_main()+
xlab("log2(dm-Grh/grh null)")+
ylab("-log10(qvalue)")+
geom_hline(yintercept = -log10(0.05), linetype = "dashed", color = "red") +
guides(colour = "none")+
theme(axis.title = element_text(size = 13))
ggsave(rna_volc, file = "./plots/rna_volcano.png", width = 3.5, height= 3)

# Percent dm-Rescue activated v repressed
rna_ce_prop_rep_act <- rna_small_direct[rna_small_direct$padj_dm < 0.05 & rna_small_direct$direct_target == "YES",] %>%
ggplot(aes(x=log2FoldChange_dm < 0, y=1-abs(log2FoldChange_ce-log2FoldChange_dm)/abs(log2FoldChange_dm)))+
geom_boxplot(notch=TRUE)+
theme_main()+
xlab("")+
ylab("Proportion of dm-Grh rescue")+
scale_x_discrete(labels=c('Grh-activated', 'Grh-repressed'))+
ylim(0,1)
ggsave(rna_ce_prop_rep_act, file = "./plots/rna_ce_prop_rep_act.png")


#################################################################
########## Chimeras and point mutations -- Figure 6 #############
#################################################################
# Rescue plots
## dm/ce
atac_rescue_fw <- atac_grh[atac_grh$padj_dm < 0.05 & atac_grh$grh_overlap == "YES",] %>%
ggplot(aes(x=log2FoldChange_dm, y=log2FoldChange_fw))+
geom_point(alpha=0.7, size=4.5)+
ylim(-0.3,1.7)+
xlim(-0.3,1.7)+
geom_hline(yintercept = 0, linetype = "dashed") +
geom_vline(xintercept = 0, linetype = "dashed") +
geom_abline(intercept = 0, slope = 1, linetype = "dashed") + theme_main() + geom_vline(xintercept = 0, linetype = "dashed")+
ylab("dm/ce-Grh effect on accessibility\nlog2(dm/ce-Grh/grh null)")+
xlab("dm-Grh effect on accessibility\nlog2(dm-Grh/grh null)")
ggsave(atac_rescue_fw, file = "./plots/ce_rescue_ATAC_small_chi_dmce.png", height = 5.5, width = 6)
## ce/dm
atac_rescue_wf <- atac_grh[atac_grh$padj_dm < 0.05 & atac_grh$grh_overlap == "YES",] %>%
ggplot(aes(x=log2FoldChange_dm, y=log2FoldChange_wf))+
geom_point(alpha=0.7, size=4.5)+
ylim(-0.3,1.7)+
xlim(-0.3,1.7)+
geom_hline(yintercept = 0, linetype = "dashed") +
geom_vline(xintercept = 0, linetype = "dashed") +
geom_abline(intercept = 0, slope = 1, linetype = "dashed") + theme_main() + geom_vline(xintercept = 0, linetype = "dashed")+
ylab("dm/ce-Grh effect on accessibility\nlog2(ce/dm-Grh/grh null)")+
xlab("dm-Grh effect on accessibility\nlog2(dm-Grh/grh null)")
ggsave(atac_rescue_wf, file = "./plots/ce_rescue_ATAC_small_chi_cedm.png", height = 5.5, width = 6)
## dm-ItoA
atac_rescue_ia <- atac_grh[atac_grh$padj_dm < 0.05 & atac_grh$grh_overlap == "YES",] %>%
ggplot(aes(x=log2FoldChange_dm, y=log2FoldChange_ia))+
geom_point(alpha=0.7, size=4.5)+
ylim(-0.3,1.7)+
xlim(-0.3,1.7)+
geom_hline(yintercept = 0, linetype = "dashed") +
geom_vline(xintercept = 0, linetype = "dashed") +
geom_abline(intercept = 0, slope = 1, linetype = "dashed") + theme_main() + geom_vline(xintercept = 0, linetype = "dashed")+
ylab("dm/ce-Grh effect on accessibility\nlog2(dm-ItoA-Grh/grh null)")+
xlab("dm-Grh effect on accessibility\nlog2(dm-Grh/grh null)")
ggsave(atac_rescue_ia, file = "./plots/ce_rescue_ATAC_small_ia.png", height = 5.5, width = 6)
## all
atac_grh_melt <- melt(atac_grh, id.vars = "gene", meas.vars = c("log2FoldChange_dm", "log2FoldChange_ce", "log2FoldChange_fw", "log2FoldChange_wf", "log2FoldChange_ia"))

a <- atac_grh[atac_grh$padj_dm < 0.05 & atac_grh$grh_overlap == "YES",] %>%
ggplot()+
#geom_point(aes(x=log2FoldChange_dm, y=log2FoldChange_ia),alpha=0.7, size=4.5, color="steelblue3")+
geom_point(aes(x=log2FoldChange_dm, y=log2FoldChange_ce),alpha=0.7, size=4.5, color="red3")+
#geom_point(aes(x=log2FoldChange_dm, y=log2FoldChange_fw),alpha=0.7, size=4.5, color="green3")+
#geom_point(aes(x=log2FoldChange_dm, y=log2FoldChange_wf),alpha=0.7, size=4.5, color="purple3")+
ylim(-0.3,1.7)+
xlim(-0.3,1.7)+
geom_hline(yintercept = 0, linetype = "dashed") +
geom_vline(xintercept = 0, linetype = "dashed") +
geom_abline(intercept = 0, slope = 1, linetype = "dashed") + theme_main() + geom_vline(xintercept = 0, linetype = "dashed")+
ylab("dm/ce-Grh effect on accessibility\nlog2(dm-ItoA-Grh/grh null)")+
xlab("dm-Grh effect on accessibility\nlog2(dm-Grh/grh null)")
ggsave(a, file = "./plots/ce_rescue_ATAC_color_ce.png", height = 5.5, width = 6)

b <- atac_grh[atac_grh$padj_dm < 0.05 & atac_grh$grh_overlap == "YES",] %>%
ggplot()+
#geom_point(aes(x=log2FoldChange_dm, y=log2FoldChange_ia),alpha=0.7, size=4.5, color="steelblue3")+
geom_point(aes(x=log2FoldChange_dm, y=log2FoldChange_ce),alpha=0.15, size=4.5, color="red3")+
geom_point(aes(x=log2FoldChange_dm, y=log2FoldChange_fw),alpha=0.7, size=4.5, color="green3")+
#geom_point(aes(x=log2FoldChange_dm, y=log2FoldChange_wf),alpha=0.7, size=4.5, color="purple3")+
ylim(-0.3,1.7)+
xlim(-0.3,1.7)+
geom_hline(yintercept = 0, linetype = "dashed") +
geom_vline(xintercept = 0, linetype = "dashed") +
geom_abline(intercept = 0, slope = 1, linetype = "dashed") + theme_main() + geom_vline(xintercept = 0, linetype = "dashed")+
ylab("dm/ce-Grh effect on accessibility\nlog2(dm-ItoA-Grh/grh null)")+
xlab("dm-Grh effect on accessibility\nlog2(dm-Grh/grh null)")
ggsave(b, file = "./plots/ce_rescue_ATAC_color_ce_fw.png", height = 5.5, width = 6)

c <- atac_grh[atac_grh$padj_dm < 0.05 & atac_grh$grh_overlap == "YES",] %>%
ggplot()+
#geom_point(aes(x=log2FoldChange_dm, y=log2FoldChange_ia),alpha=0.7, size=4.5, color="purple3")+
geom_point(aes(x=log2FoldChange_dm, y=log2FoldChange_ce),alpha=0.15, size=4.5, color="red3")+
#geom_point(aes(x=log2FoldChange_dm, y=log2FoldChange_fw),alpha=0.15, size=4.5, color="green3")+
geom_point(aes(x=log2FoldChange_dm, y=log2FoldChange_wf),alpha=0.7, size=4.5, color="steelblue3")+
ylim(-0.3,1.7)+
xlim(-0.3,1.7)+
geom_hline(yintercept = 0, linetype = "dashed") +
geom_vline(xintercept = 0, linetype = "dashed") +
geom_abline(intercept = 0, slope = 1, linetype = "dashed") + theme_main() + geom_vline(xintercept = 0, linetype = "dashed")+
ylab("dm/ce-Grh effect on accessibility\nlog2(dm-ItoA-Grh/grh null)")+
xlab("dm-Grh effect on accessibility\nlog2(dm-Grh/grh null)")
ggsave(c, file = "./plots/ce_rescue_ATAC_color_ce_wf.png", height = 5.5, width = 6)

d <- atac_grh[atac_grh$padj_dm < 0.05 & atac_grh$grh_overlap == "YES",] %>%
ggplot()+
geom_point(aes(x=log2FoldChange_dm, y=log2FoldChange_ia),alpha=0.7, size=4.5, color="purple3")+
geom_point(aes(x=log2FoldChange_dm, y=log2FoldChange_ce),alpha=0.15, size=4.5, color="red3")+
#geom_point(aes(x=log2FoldChange_dm, y=log2FoldChange_fw),alpha=0.15, size=4.5, color="green3")+
#geom_point(aes(x=log2FoldChange_dm, y=log2FoldChange_wf),alpha=0.7, size=4.5, color="steelblue3")+
ylim(-0.3,1.7)+
xlim(-0.3,1.7)+
geom_hline(yintercept = 0, linetype = "dashed") +
geom_vline(xintercept = 0, linetype = "dashed") +
geom_abline(intercept = 0, slope = 1, linetype = "dashed") + theme_main() + geom_vline(xintercept = 0, linetype = "dashed")+
ylab("dm/ce-Grh effect on accessibility\nlog2(dm-ItoA-Grh/grh null)")+
xlab("dm-Grh effect on accessibility\nlog2(dm-Grh/grh null)")
ggsave(d, file = "./plots/ce_rescue_ATAC_color_ce_ia.png", height = 5.5, width = 6)
```
